# Growth consequences of the inhomogeneous organization of the bacterial cytoplasm

**DOI:** 10.1101/2023.04.18.537336

**Authors:** Johan H. van Heerden, Alicia Berkvens, Daan H. de Groot, Frank J. Bruggeman

**Affiliations:** Systems Biology Lab, Amsterdam Institute for Molecules, Medicines and Systems (AIMMS), VU Amsterdam, De Boelelaan 1085, NL-1081 HV Amsterdam, The Netherlands; Computational Systems Biology, Biozentrum, Universitaet Basel

**Keywords:** *Escherichia coli*, fluorescence microscopy, mathematical model, single cell growth rate, balanced growth, inhomogeneous cytoplasm, nucleoid exclusion, cell cycle dynamics

## Abstract

In many bacteria, translating ribosomes are excluded from the nucleoid, while amino-acid and energy-supplying metabolic enzymes spread evenly throughout the cytoplasm. Here we show with time-lapse fluorescence microscopy that this inhomogeneous organisation of the cytoplasm can cause single *Escherichia coli* cells to experience an imbalance between biosynthesis and metabolism when they divide, resulting in cell size-dependent growth rate perturbations. After division, specific growth rate and ribosome concentration correlates negatively with birth-size, and positively with each other. These deviations are compensated during the cell-cycle, but smaller-than-average cells do so with qualitatively different dynamics than larger-than-average cells. A mathematical model of cell growth, division and regulation of biosynthetic and metabolic resource allocation reproduces our experimental findings, suggesting a simple mechanism through which long-term growth rate homeostasis is maintained while heterogeneity is continuously generated. This work shows that the life of single bacterial cells is intrinsically out-of-steady-state, dynamic and reliant on cytoplasmic organization.

**Popular summary:** Classical, population-level studies of the metabolism and growth of bacteria indicate that the average cell in a growing population operates at steady state and can be viewed as an homogeneous ‘bag of enzymes’. Here we show that this view does not capture the lives of single cells. At birth, they are perturbed from the steady state of their mother cell after which they need their entire cell cycle to return to this state by active regulation. Then they divide and their daughters are perturbed again; a never ending cycle that is inescapable and akin to a Sisyphean task. This behaviour emerges from the delicate interplay of the intrinsic randomness of (uneven) cell division, the inhomogeneous localisation of metabolic and ribosomal proteins in the cell, unbalanced metabolism, and compensatory steering of gene expression.

## Introduction

When cultivation conditions are held constant, isogenic bacterial populations eventually converge to a state of balanced growth, characterised by a time-invariant per-capita growth rate, and constant properties of the ‘average cell’, which has a metabolism operating at steady state. ^1–3^ Under such conditions, population growth is stationary and the fraction of cells at a specific point in the cell cycle remains constant. Moreover, the average cell is then generally considered ‘ideally stirred’, lacking any spatiotemporal organisation. This growth condition enabled quantitative bacterial physiology ^4,5^ and is the reference state for most contemporary systems-biological studies on metabolism and growth. ^1,3,6^

Since single cells display inevitable, random fluctuations (noise) of molecule concentrations, cellsize and growth rate ^1,7–9^, population characteristics can only remain time invariant, when noise is either compensated for ^10^ or remains negligible. That noise in concentrations of abundant proteins is likely negligible is captured by the rule of thumb that it is proportional to 1 over the mean copy number. ^11^ This leads to the prediction that fluctuations in abundant proteins associated with biosynthesis are generally insignificant. However, fluctuations in cellular growth rate have been found ^8^, suggesting the existence of unknown systemic origins of noise. ^12,13^ On top of that, single cells fluctuate in birth and division size during balanced growth. The mechanisms underlying size homeostasis of single cells have received considerable attention in recent years ^1,10,14–19^, while the variability and homeostasis of single-cell growth rate remain poorly understood.

It is becoming increasingly clear that birth-size dependent deviations from exponential growth occur along bacterial cell cycles. ^17,20,21^ Thus, single cells can deviate from the balanced growth state of the average cell in a systematic manner, challenging the concept that single cells are growing exponentially in size along their cell cycle, like the average cell and cell populations do during balanced growth. The origins of systematic cell cycle-dependent growth rate deviations are, however, not understood, nor is it known how widespread this phenomenon is. These aspects we address in this paper.

At balanced growth of symmetrically dividing cells, such as *E. coli* and *B. subtilis*, the size and molecular content of the average cell doubles during each cell cycle and is halved at cell division. ^7^ In single cells, the underlying processes are subject to random fluctuations that, amongst other effects, influence the precision of cell division into equally sized halves and the partitioning of different cellular components (e.g. (macro)molecules) into newborn cells. ^7^ It is well known that low copynumber components, like transcription factors and plasmids, are prone to partitioning errors, but these tend to be random ^22,23^, incapable of giving rise to systematic deviations. On the other hand, when high copy number and homogeneously dispersed components are partitioned, concentrations will generally be insensitive to imperfect cell division. ^23^ Size differences between newborn cells will, however, impact concentrations of spatially-localised cellular components. ^23^

The bacterial cytoplasm displays spatial organisation. ^24–28^ For example, protein aggregates or large assemblies (e.g. ribosomes) preferentially localise to specific cellular regions and are expelled from the nucleoid. ^26,29–31^ For instance, in rod-shaped bacteria such as *E. coli* and *B. subtilis*, fully-assembled, translating ribosomes are excluded from the mid-cell positioned nucleoid and confine predominantly to cell poles. ^25–28,32,33^ Simulations indicate that this phenomenon likely results from volume exclusion forces (an entropic effect). ^25,26,30,34^ Smaller (macro)molecules, such as metabolic proteins and their reactants, move freely through the nucleoid mesh, dispersing homogeneously throughout the cytoplasm. ^25,26^

We speculated that the different localisation patterns of ribosomes and metabolic proteins may cause an imbalance in the metabolism of newborn cells, due to a perturbation of their relative concentrations after an uneven cell-division event. We hypothesised that this can result in a growthrate perturbation at cell birth. This effect likely correlates with birth size since smaller-than-average newborn cells have a relative higher polar volume fraction (where the majority of ribosomes reside) than average cells, while larger-than-average cells have a lower fraction. This would explain recent observations of structured size-dependent growth-rate perturbation of newborn cells ^17,20,21^. Also, since ribosomes are excluded from the nucleoids of many bacterial species, ^25,27^ this mechanism may be widespread.

Here we tested these hypotheses. We studied how growth-rate variability and homeostasis in single *E. coli* cells is influenced by cell division and compensatory processes along the cell cycle. Our results confirm that imperfect cell division, the localisation of ribosomes, and homogeneously disersed metabolic proteins results in imbalances in metabolic and ribosomal protein concentrations in newborn cells. This imbalance causes smaller-than-average cells to grow faster than average-sized cells at birth, while larger-than-average cells grow slower. We present a generic mathematical model of cell growth, incorporating spatially-localised ribosomes and regulation of biosynthetic resource allocation, giving rise to growth-rate perturbations at cell birth and restoration of a balanced growth-rate at cell division. Simulations qualitatively capture size-dependent growth rate dynamics of single *E*.*coli* cells growing on defined and rich media, respectively. Our findings suggest that the spatial organisation of the bacterial cytoplasm has a significant impact on cellular heterogeneity and can disturb cellular homeostasis and growth, necessitating compensatory regulation. Furthermore, this work highlights a novel kind of systematic cell-to-cell heterogeneity in bacterial populations that grow balanced, and reinforces the question whether population averages ever truly reflect the state of individual cells.

## Results

### The growth-rate dynamics of single cells along their cell cycles depends on their birth size

We monitored single *Escherichia coli* cells during balanced growth, using live-cell imaging with fluorescence microscopy. We validated balanced growth by confirming the time invariance of the probability distribution of cell ages, sizes, generation times and other key properties (Fig. S1), ^35^ to ensure that cells with an equal cell-cycle progression can be compared, even though they were observed at different times ^20^. This allows us to determine the dynamics of the specific growth rate of cell length (dlnL/dt), which we call the specific elongation rate (sER), as function of cell-cycle progression (the normalised cell age) (Fig. 1). Throughout, when referring to single-cell behaviours, growth rate implies elongation rate. The cell-cycle progression of a single cell is defined as the ratio of the elapsed time since birth over its generation time, such that cells are born at normalised age 0 and divide at normalised age 1.

**Figure 1:**
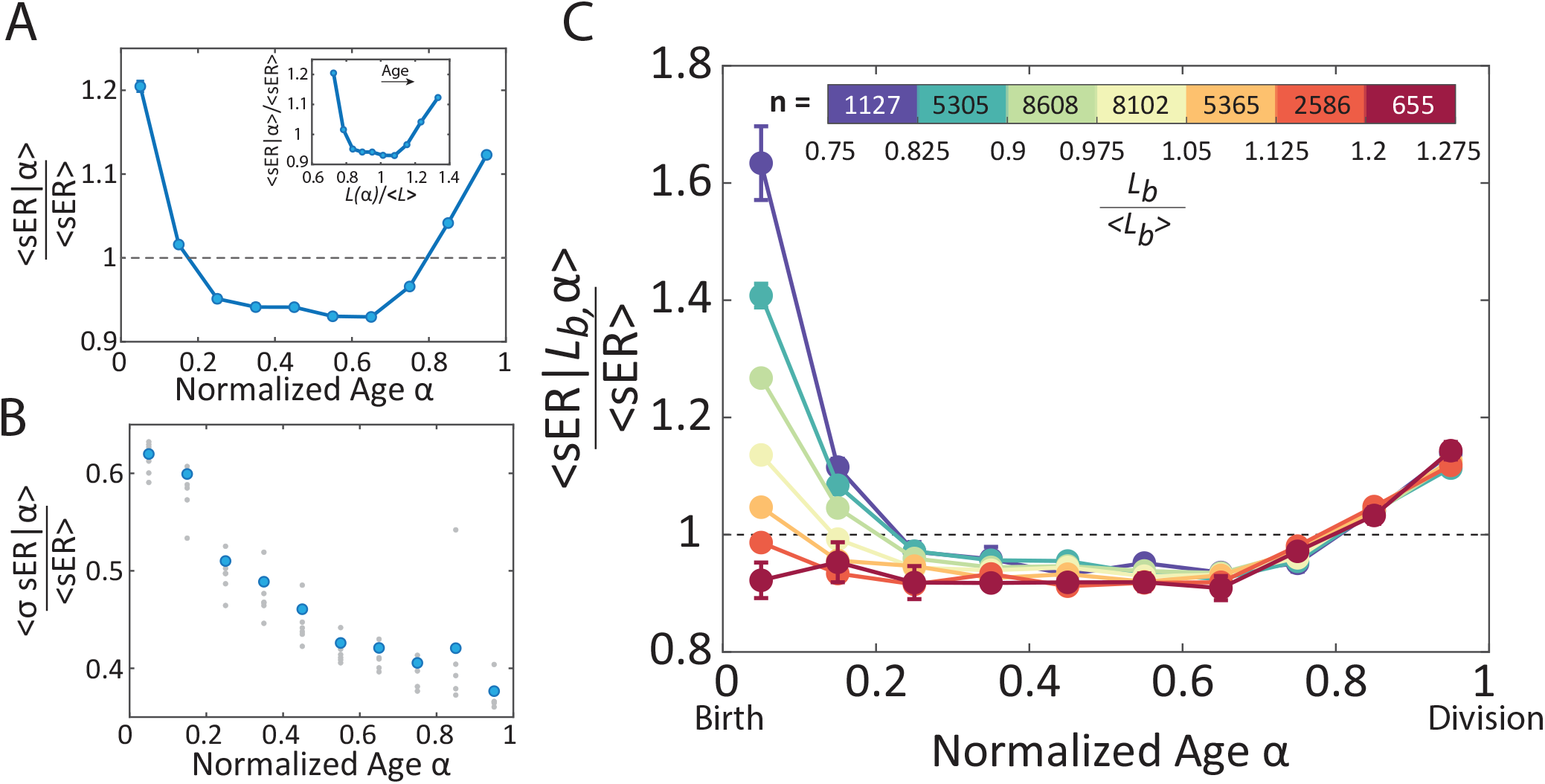
Systematic deviations and the birth-size dependence of the specific elongation rate as function of cell cycle progression. Shown is data for *E. coli* cells growing on glucose. We denote the specific elongation rate by *sER* and an average of a variable *x* is denoted by (*x*). (A) The average specific elongation rate (*sER*) of single cells changes as function of their normalized cell age (age 0 corresponds to cell birth and age 1 to division). Exponential growth corresponds to the horizontal, dashed line. The inset figure shows this same trend, but as a function of the average length per normalized age, normalized by the total average length. Since the average length observed in a growing population corresponds to a cell approximately half way through its cell cycle, we note that the expected range runs from 2/3 to 4/3. (B) The coefficient of variation of the sER of single cells decreases as a function of normalized age, indicating compensatory dynamics. (Blue markers indicate the average of the biological replicates shown as grey markers.) (C) The mean sER of different birth-length bins are shown as function of the normalized cell age. The data indicates that smaller-than-average cells (dark blue) grow faster than average-sized cells (light green) while larger-than-average cells (red) grow slightly slower. The bar legend indicates the normalised ranges of the birth-length bins, and the number of individual cells included in each bin. All plots show the average from 7 independent experiments, all for cells grown on defined minimal medium (M9) with glucose as carbon source and with a total of 31,748 cells. Error bars are plotted as standard error of the mean; where they are not visible they are smaller than the plot markers.

Figure 1 indicates that the specific growth rate of the average single cell (averaged across 31,748 individual cells) displays systematic deviations along the cell cycle; a finding that corresponds to earlier observations with *Bacillus subtilis* ^20,21^. At cell birth, the sER is higher than the cell-cycle average, after which it decreases, becomes approximately constant, and finally rises towards the end of the cell cycle (Fig. 1A). Additionally, variability of elongation rates of individual cells (quantified by the standard deviation) is highest at birth, after which it declines rapidly as cells proceed through their cell cycle (Fig. 1B). These observations indicate that cell division induces growth rate differences between individual newborn cells that are subsequently compensated during the cell cycle.

To investigate the origin of this growth rate perturbation at birth, we divided the growth rate data into classes (bins) of cells with similar birth sizes (see Table S1 for details). A striking pattern is then observed: birth size correlates negatively with the growth rate (sER) at birth (Fig. 1C). This correlation was also observed in *B. subtilis* ^20^. It indicates that smaller-than-average newborn cells grow significantly faster than average newborn cells, while larger-than-average newborn cells grow slower, in line with recent findings by Vashistha et al. ^21^. These growth-rate differences rapidly decrease as cells progress through their cell cycle (Fig. 1C). By the end of the cell cycle, the growth rate (sER) has become independent of birth size and indistinguishable from the rate of the average cell (Fig. 1C). These findings show that growth-rate fluctuations correlate with size perturbations at cell birth and suggest an active compensation of the growth rate perturbation along the cell cycle. In addition, non-average sized newborns have partially corrected their length deviation at the end of the cell cycle, in agreement with observations of *E. coli* by Wallden et al. ^16^ (Fig. S3 and S5).

### The ribosome concentration of newborn cells correlates with their size and growth rate

Figure 2A shows the dependency of the growth rate at birth on birth size. It shows that smallerthan-average cells grow faster than average cells and larger-than-average cells grow slower.

**Figure 2:**
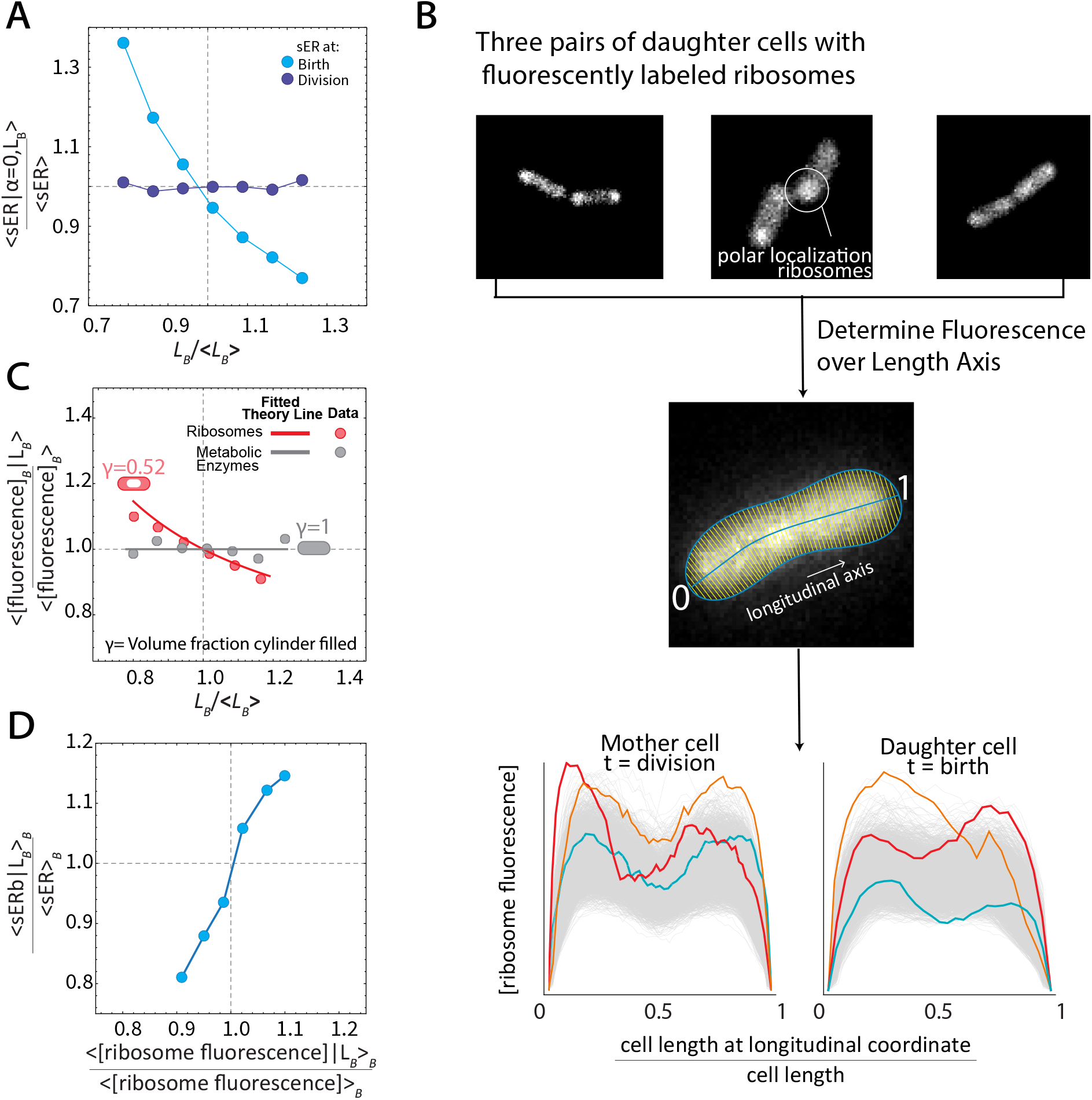
The mean growth rate and the ribosome concentration of newborn cells correlates negatively with their size. (A) The normalized specific growth rate of newborn cells is shown as function of the normalised birth size. Normalisation was done by division by the corresponding average value. (B) Three pairs of daughter cells are shown with fluorescently-labelled ribosomes that preferentially localise outside of the nucleoid in cell poles (Top). We quantified the ribosome concentration along the length axis (at fixed length intervals) of single cells by determining the total fluorescence orthogonal to this axis (Middle). The resulting data is shown as function of the normalised cell length (Bottom), which indicates the highest ribosome fluorescence at cell poles. Note that at the cell ends, the fluorescence drops in the periplasmic region of cells, which is devoid of ribosomes. (C) The concentration (total fluorescence divided by cell area) of the ribosome at birth and GFP are shown as function of the birth size of newborn cells, indicating that smaller-than-average cells have indeed higher ribosome concentrations than average-sized whereas larger-than-average cells have lower concentrations. The concentration of GFP proteins in newborn cells is size independent, because they spread homogeneously, like metabolic proteins. The full lines are fits of theoretical expectations for a cell that is either completely filled with a homogeneously spread protein (*γ*=1) or with ribosomes that occupy about half of the cytosolic volume (*γ*=0.52). For the GFP protein data, cells were grown on lactose as carbon source. (D) The normalised mean growth rate of newborn cells is plotted against their ribosome content (as function of their birth size; results of Figure A and C combined), showing that the growth rate of newborn cells correlates positively with their ribosome content, as expected ^6^.

Since the growth rate is constant at metabolic steady state ^3^, we speculated that cell birth induces a metabolic imbalance that is dependent on cell size. We tested whether this imbalance is caused by a size-dependent perturbation in the ribosome concentration of newborn cells, as growth rate is proportional to the ribosome concentration. ^6^

We used a previously validated strain ^32^ with an mCherry-tagged L9 ribosomal subunit to correlate the growth rate of a newborn cell and its ribosomal content with its (birth) size (Fig. 2B). Our expectation that the ribosome concentration of a newborn cell depends on its birth size, stems from the spatial organisation of the cytoplasm of many bacterial cells. ^27^ Rod-shaped bacteria such as *E. coli* have a mid-cell positioned nucleoid ^27,28^ from which ribosomes are excluded, due to their large size. ^30^ This exclusion leads to enrichment of ribosomes in the cell poles. ^27,32,33^ Our data indeed confirms this (Fig. 2B). We note that RNA polymerase complexes localise in the nucleoid ^33^ and that small metabolic proteins are homogeneously spread throughout the cytoplasm. ^26^ These proteins are not excluded, because they are smaller than ribosomes, and small enough to diffuse freely through the nucleoid mesh. ^25,26,30^

If we assume (for simplicity) that all ribosomes are localised in cell poles, we propose a mechanism of partitioning that works as follows (Fig. 3A; we present the general case in the SI): When a mother cell divides, daughter cells receive the same number of ribosomes, regardless of their cell volume, because they each obtain two equally-sized poles, and ribosome copy-number fluctuations are negligible due to their high average values (up to several tens of thousands per cell ^26,33^). In contrast, the number of metabolic proteins they receive is proportional to their cell volume (since they are spread homogeneously). Smaller-than-average newborn cells have a higher polar-volume fraction and will therefore have an excess of ribosomes (relative to metabolic proteins) compared to the average cell and grow faster at birth (because growth rate is proportional to the ribosome concentration ^6^), while large newborn cells have a lower polar-volume fraction than average cells, and therefore a proportional ribosome shortage such that they grow slower at birth. This is indeed what we found when we correlated the cell length of newborn cells with their ribosome concentration (Fig. 2D).

**Figure 3:**
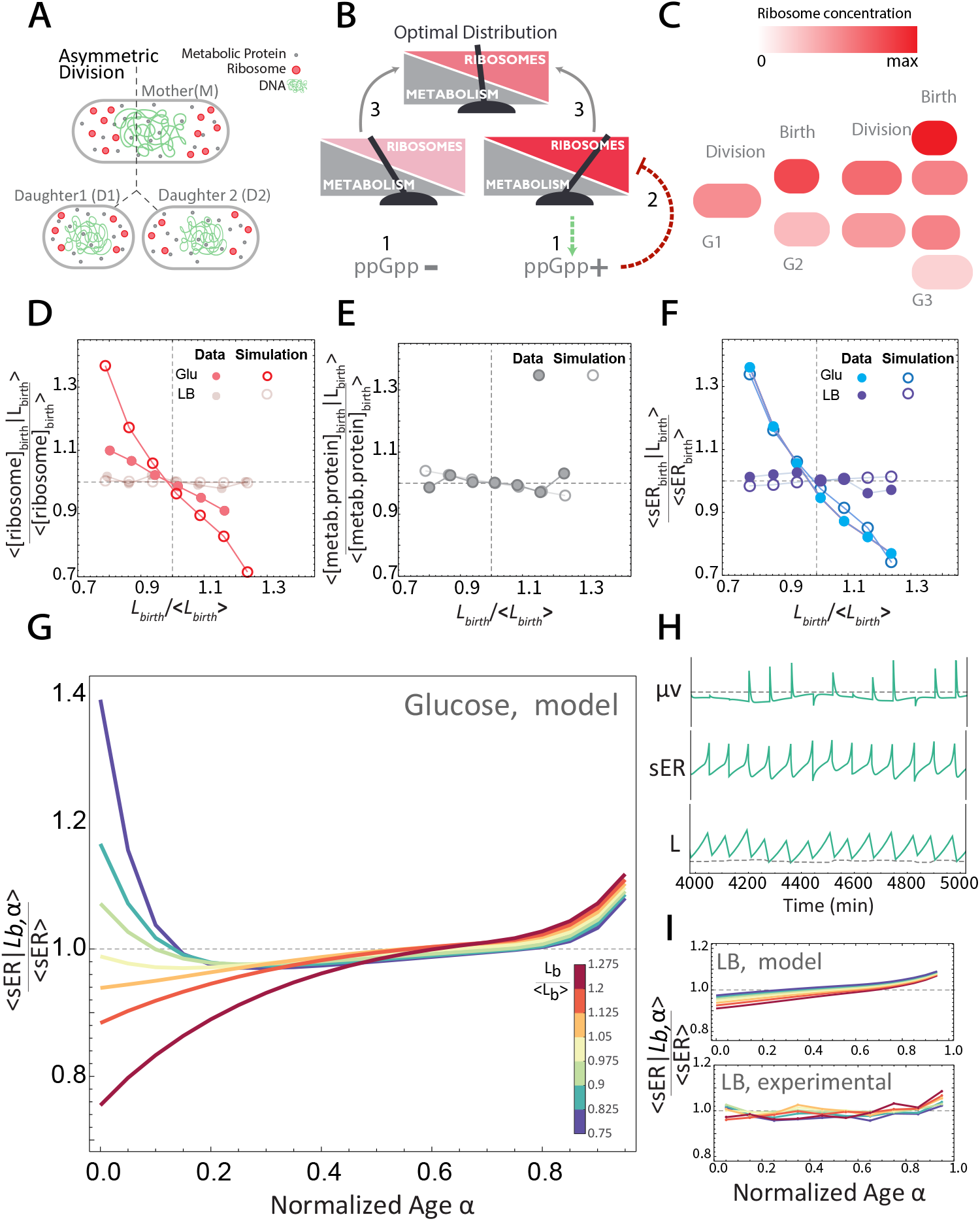
Generic mathematical model captures the experimental data. (A) Uneven division of a mother cell with polar ribosomes and homogeneously spread metabolic enzymes leads to daughter cells with identical metabolic enzyme concentrations, but deviating ribosome concentrations. Small newborn cells tend to have higher ribosome concentrations than large newborn cells. (B) In the model, ppGpp regulates the ratio of ribosomes to metabolic enzymes, by steering the saturation of ribosomes with their substrates to a fixed setpoint. (C) Since restoring the optimal metabolism versus biosynthesis rates (as explained in B) takes more than one cell cycle, mother cells will generally not yet be in a steady state at their division. Their daughters therefore inherit the perturbation consequences of previous generations, affecting their growth rate at birth. (D-F) A comparison of the experimental data for differently sized cells at birth, to averages of 50000 consecutive cell-cycle simulations. For ribosomes in D, metabolic enzymes in E and sER in F. (G) Birth-to-division growth rate trajectories for different length bins, from our mathematical model simulation with 50000 rod-shaped cells, based on cell-size dependent ribosome partitioning (A), saturation set-point control of ribosome expression (B), and the non-steady state mother effect (C). This figure qualitatively captures the experimental data shown in Fig. 1C. (H) A panel of three plots showing representative simulated trajectories of volumetric growth rate (*μ*_*V*_), elongation rate and cell length. For the growth rate plot, the dashed line indicates the average growth rate for the entire simulation. For the length plot, the dashed line indicates 50% of the length of the associated mother cell. (I) Comparison and validation of experimental data with a model prediction. The growth-rate effect of polar localisation of ribosomes is less in large cells, because a relatively large fraction of ribosomal is located mid-cell, along the nucleoid ^33,45^, which reduces the size-dependent asymmetry in ribosome and metabolic protein concentration in non-average-sized, newborn cells.

In contrast to the ribosome partitioning pattern, a constitutively expressed and homogeneously distributed fluorescent protein (GFP), mimicking a small (metabolic) protein, shows no concentration relation with birth size (Fig. 2C). The measured relationship in Figure 2C matches the expected theoretical relationship of a rod-shaped cell (see SI) with 48% of its cytoplasm occupied by the nucleoid, which is a realistic value (cf. 46-75% ^33,36,37^). We note that daughter cells of unevenly divided mothers will generally have different ribosome concentrations and as a result grow at different rates.

We conclude that the asymmetric spatial localisation of ribosomes and metabolic enzymes causes a catabolic-anabolic imbalance at cell division that leads to a size-dependent variability of the growth rate of newborn cells. In the next section, we attempt to explain how single cells achieve compensation of the growth-rate disturbance at birth, to eventually attain an (almost) size-independent growth rate prior to the next division.

### Compensatory regulation can correct the metabolism-biosynthesis imbalance of newborn cells and restore growth-rate homeostasis at the end of the cell cycle

Figure 2 confirmed that on average smaller-than-average newborn cells will be confronted with a relative excess of ribosomes over metabolic enzymes, while larger-than-average cells experience a shortage. We expect that this results in a size-dependent, transient imbalance between synthesis (metabolic) and consumption (ribosomal) of amino acids. In smaller-than-average cells, the immediate effect of a ribosome excess is a rise in the growth rate, since growth rate is proportional to protein synthesis rate which is proportional to ribosome content. This enhancement of the growth rate diminishes over time, because of the depletion of amino acids, as the metabolic protein concentration of these cells is too low to keep up with the high amino acid demand of the excess ribosomes. In the larger-than-average cell, the reverse happens: the relative shortage of ribosomes reduces growth rate at birth. But, since these cells have excess metabolic proteins over ribosomes (so a relative ribosome shortage), the amino-acid supply rate exceeds the demand by ribosomes. This imbalance results in the rise of the amino-acid concentration in these cells and an increase in the protein-synthesis rate of ribosomes, leading, in turn, to an increase in growth rate. All of this happens on a seconds to minute (metabolic) time scale in single cells. Adjustment of protein concentrations occurs next on a slower time scale.

The transient metabolic imbalances described here also occur during nutrient up- and downshifts. In these cases, ppGpp-mediated regulatory machinery adjusts metabolic and ribosomal gene expression, such that a balance is restored ^38–44^.We propose that the same mechanism operates to restore imbalances that arise from transient internal fluctuations during growth and division cycles. In response to a relative excess of ribosomes (and amino acid shortage), ppGpp is known to rise and bind to RNA polymerase, lowering its affinity for ribosomal promoters, which leads to its enhanced allocation to catabolic promoters ^38^. This increases synthesis of metabolic proteins (at the expense of ribosomes) and a restoration of the growth rate homeostasis at the end of the cell cycle. In the larger-than-average newborn cell the opposite happens: more ribosomes are made to counter the imbalance between amino acid synthesis (metabolism) and consumption (ribosomes), thereby also restoring growth rate homeostasis. In both of these scenarios, the specific growth rate of cells with deviating newborn sizes becomes size independent at the end of the cell cycle (Fig. 1C).

Although our data (Fig. 1C) indicates that the growth rate at the end of the cell cycle becomes birth-size independent, the ribosome concentration is still dependent on birth size (Fig. S4). This indicates that restoration of a steady-state metabolism appears to take on average longer than a single cell cycle. Accordingly, the relaxation time for ribosome concentration homeostasis exceeds the generation time such that most newborn cells stem from mothers that have not yet fully compensated for ribosome deviations. This adds an additional dynamics that influences the metabolic imbalance of newborn cells, and suggests that two equally sized newborn cells can show distinct growth rates and different ribosome concentrations depending on the metabolic state of their dividing mothers.

Moreover, since both length (Fig. S3) and ribosomes (Fig. S4) deviations are not fully restored after a single cell cycle, we expect ancestral influences to have a characteristic effect. When cell width stays constant, the polar caps of smallerand larger cells will be of equal size. Then their cell-length difference is only determined by their mid-cell length such that the polar caps take up a larger volume fraction in smaller cells than in large cells (Fig. S9). We therefore expect a larger effect of uneven cell division for cells born from small mothers, as the percentage of the mother cell filled with ribosomes is larger. This birth-size asymmetry we indeed observe in the growth rates of newborn cells in our data (Fig. 1C), small cells deviate more from average than large cells.

### A mathematical model reproduces the experimental data

To test whether the above described effects of uneven division, nucleoid-excluded ribosomes, cell growth, the ppGpp-mediated regulation mechanism, and the ‘ancestral, mother effect’ (Fig. 3A-C) can indeed account for the dynamics observed in the experimental data, we developed a generic mathematical model. The model applies to rod-shaped bacteria that aim to use their ribosomes at optimal efficiency across conditions. These bacteria would then display a linear relation between their ribosomal protein fraction and growth rate ^41,46^, which is valid for *E. coli* except at very low growth rates. It has been suggested that the control objective of ppGpp-mediated regulation of ribosome expression is to maintain the saturation degree of ribosomes constant, which leads to robust, close-to-optimal ribosome expression ^41,46^ by optimal distribution of RNA polymerase over catabolic and ribosomal operons. ^38^

Our model simulates sequential cell cycles, during which a rod-shaped cell grows from birth to division, after which it divides. Birth sizes are sampled in agreement with our experimental data and conditional on the division length of the associated mother cell (Fig. S12). Uneven division of a mother cell will cause a birth-length dependent perturbation of the ribosome concentration in the newborn cell (SI eq. 11), while its metabolic protein concentration equals that of its mother. These perturbations are compensated for by a ppGpp-regulated mechanism that modulates metabolic and ribosome expression to steer the substrate-saturation of ribosomes to a desired set-point level. ^41,46^ Thousands of sequential cell cycles were simulated (Fig. 3H) to capture the effect that cells are typically not born from mothers in a homeostatic state, but inherit instead the effects of perturbations of previous generations (Fig. 3C). This model is capable of qualitatively reproducing the compensatory dynamics of the growth-rate perturbation that we observe in the experimental data of *E. coli* (Fig. 1C and 3G).

The experimental data (Fig. 1C) and the model simulations (Fig. 3G) both show an increase in the specific elongation rate of cells close to the end of the cell cycle. This is most likely due to the invagination of the cell wall during septum formation and cell growth at a constant volumetric growth rate such that the length growth rate increases when the diameter of the constricting, mid-cell region becomes smaller to eventually approaches zero, after which division occurs (Fig. S10).

### Experimental confirmation of a model prediction

Our model makes a testable prediction about the behaviour of long, rod-shaped cells. It predicts that the magnitude of the growth rate perturbation at birth depends on the fraction of the ribosomes found in cell poles versus those that surround the nucleoid in the non-polar, mid-cell region of the cell. In a long, rod-shaped cell, where the polar volume is only a small portion of the total cell volume, the ribosomes are still excluded from the nucleoid, but many of them now reside along side the nucleoid, in the mid-cell region. The model predicts that under conditions when cells are long, uneven division perturbs the ribosome concentration in non-averaged-sized newborn cells less than in cells that are grown under conditions when the average cell is smaller (and the polar volume fraction is larger) (Fig. 3I vs G). Thus, we would not expect to see similar significant size-dependent growth-rate perturbations at cell birth when cells are grown in conditions where they are large on average.

To test this, we grew *E. coli* on a complex rich medium (Luria broth; LB) where the average cell length is 2 times larger and cells are 1.5 times wider than on a minimal medium with glucose as a carbon source (Tables S1-S3). Indeed, we find now that both the growth rate (Fig. 3F and 3I) and the ribosome concentration at birth (Fig. 3D and S4B) are no longer size dependent, with relations resembling those seen for a homogeneously dispersed protein (Fig. 2C, 3E and S4B and C). This behaviour we can reproduce in the model by keeping all parameters the same, except for the division length of the cell, the relative time of septum cap formation ^16^ and fraction of the cylindrical cytoplasm filled with ribosomes.

## Discussion

A defining characteristic of balanced growth is that the population averages of cell lengths at birth and division, generation times and growth rate along the cell cycle are time invariant (homeostatic) ^10^, despite random fluctuations in these quantities. As a consequence, the frequencies of small newborn cells that grow faster than average do not increase over time, nor does the frequency of larger than average cells decrease. In this manner, homeostasis of cell size (birth, average and division size) is preserved. The specific growth rate of a cell population at balanced growth is also constant. Here we showed that at a single cell level this quantity shows large perturbations at cell birth due to an imbalance in the rate of metabolism and biosynthesis, caused by an asymmetry in the localisation of ribosomes and metabolic proteins in single cells.

The work of Gray et al. ^27^ highlights the omnipresence of mid-cell positioning and compaction of bacterial nucleoids, suggesting that the phenomenon we described in this paper may be widespread among bacteria. Our work suggests that bacteria that experience exclusion of ribosomes from their nucleoid are prone to show growth-rate perturbations at cell birth. A key parameter is the fraction of ribosomes in the cell poles. If most of the ribosomes surround the mid-cell nucleoid, then the number of ribosomes that a daughter cell receives will no longer be determined by the number of ribosomes in its cell poles, but rather, will start to correlate with total cellular volume. This happens in cells that are very long with only a small fraction of their volume being polar, as we showed for cells growing on complex medium (Fig. 3I). We therefore also estimate that the cell with a pole volume fraction above a critical value, such as cocci or long rod-shaped cells, no longer display these growth-rate deviations at birth.

## Conclusion

We have shown that the spatial localisation of ribosomes and asymmetrical cell division causes large perturbations of the specific growth rate in newborn rod-shaped bacterial cells. This implies that it is unlikely that single bacterial cells exhibit true balanced growth with steady-state metabolism and a constant growth rate along their cell cycle, even though a population of them can show a constant growth rate. This highlights the importance of considering the implications of steadystate assumptions when studying single cell behaviour. Finally, it is intriguing that something as fundamental as an inevitable entropic force (leading to nucleoid-exclusion of ribosome), underlies a systematic perturbation of cells at division which then necessitates persistent, compensatory control. A steering mechanism based on efficient usage of ribosomes and metabolic enzymes, by prevention of overexpression, appears a robust strategy for restoration of a balanced growth rate. The omnipresence of nucleoid-excluded ribosomes across bacteria suggests that the mechanism we report may turn out to be widespread among bacteria.

## Acknowledgments

J.v.H. and F.B. received funding from the EraSysApp grant RobustYeast. A.B. received funding from the European Union’s Horizon 2020 research and innovation programme under the Marie Sk-lodowska-Curie grant agreement no. 713669. We thank Matt Scott for this suggestions of the mechanism explaining the acceleration of the length growth rate during cell division at a constant volumetric growth rate. We thank Pieter Rein ten Wolde for critical discussions, reading of the manuscript and providing feedback, and Joen Luirink, Matti Gralka and Tanneke den Blaauwen for reading the manuscript and providing feedback.

## Author Contributions

J.v.H. and F.B. designed the study; J.v.H. performed experiments; J.v.H. and A.B. performed analyses; A.B., D.d.G and F.J.B developed the theory. J.v.H., A.B. and F.B. drafted and wrote manuscript.

## Methods

### Terminology and abbreviations

**Table 1:**
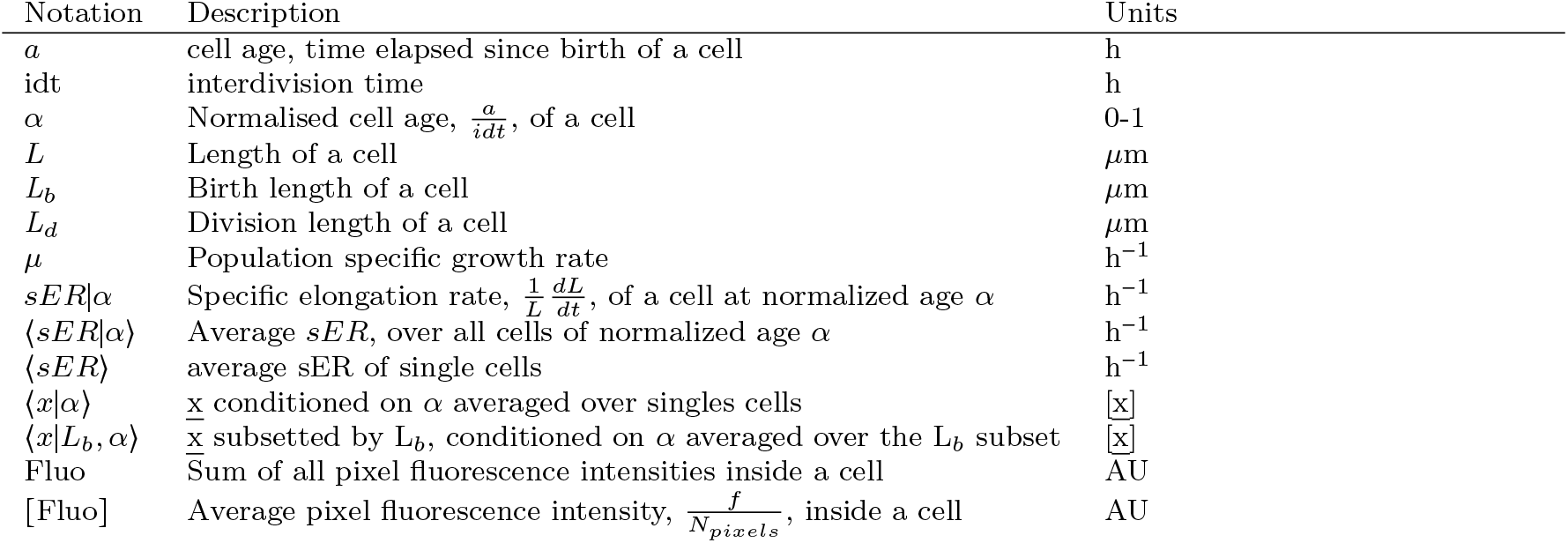
Notations used in this work.

### Strain, medium and culturing

The MG1655 derived MUK21 *Escherichia coli* strain (see ^47^ for details) was kindly provided by D. Kiviet and contains a genome integrated GFP gene under the control of the wild-type lac promoter. The MG1655 derived QC101 *E. coli* strain (see ^32^ for details) was kindly provided by S. Sanyal and contains a fusion of the red fluorescent protein mCherry with the ribosomal protein L9.

All strains were revived from glycerol stock by inoculating directly into M9 minimal medium (42.2 mM Na_2_HPO_4_, 22 mM KH_2_PO_4_, 8.5 mM NaCl, 11.3 mM (NH_4_)_2_SO_4_, 2.0 mM MgSO4, 0.1 mM CaCl_2_), supplemented with trace elements (63 *μ*M ZnSO_4_, 70 *μ*M CuCl_2_, 71 *μ*M MnSO_4_, 76 *μ*M CoCl_2_, 0.6 *μ*M FeCl_3_), 0.2 mM uracil, 1 mM Thiamine (all chemicals from Sigma) and 1 mM glucose as carbon source. At intervals of 3 hours, pre-cultures were transferred twice to fresh medium containing either M9 + 10 mM Lactose, M9 + 20 mM Glucose or LB. Additionally, 1 mM IPTG or 25 *μ*g/ml Kanamycin was included as indicated (see Table S1). After 2 transfers, an overnight culture was inoculated to a final optical density (OD, 600 nm) of ∼ 2.5 × 10^−6^ for the Glucose and Lactose experiments, and ∼ 1 × 10^−9^ for the LB experiments. After 16 hours (Glucose and Lactose) and 13 hours (LB), the cultures were again diluted to an OD600 of ∼ 2.5 × 10^−3^. Once the culture reached an OD600 of 0.01, 2 *μ*L was transferred to a 1.5% low melt agarose pad (∼ 5 mm^2^) freshly prepared with either M9 + 0.2 mM Uracil, 1 mM Thiamine and Carbon source (10 mM Lactose or 20 mM Glucose, i.e. a total of 120 C-mM) or LB, and 1 mM IPTG or 25 *μ*g/ml Kanamycin as indicated (see Table S1).

All cultures were incubated at 37 °C, in an orbital shaker at 200 rpm. Once seeded with cells, agarose pads were inverted and placed onto a glass bottom microwell dish (35 mm dish, 14 mm microwell, No. 1.5 coverglass) (Matek, USA), which was sealed with parafilm and immediately taken to the microscope for time-lapse imaging.

### Microscopy

Imaging was performed with a Nikon Ti-E inverted microscope (Nikon, Japan) equipped with 100X oil objective (Nikon, CFI Plan Apo *λ* NA 1.45 WD 0.13), Zyla 5.5 sCmos camera (Andor, UK), brightfield LED light source (CoolLED pE-100),fluorescence LED light source (Lumencor, SOLA light engine), GFP (Excitation: 460-500 nm, Dichroic: 505 nm LP, Emission: 510-560 nm) and mCherry (Excitation: 560-580 nm, Dichroic: 600 nm LP, Emission: 610 nm LP) filter sets, computer controlled shutters, automated stage and incubation chamber for temperature control. Temperature was set to 37 °C at least three hours prior to starting an experiment. Nikon NISElements AR software was used to control the microscope.

Brightfield images (80 ms exposure time at 3.2% power) were acquired every minute for Glucose and Lactose experiments and every 30 s for LB experiments. GFP fluorescence images (1 second exposure at 25% power) were acquired every 10 minutes for strain Muk21. For strain QC101, mCherry fluorescence images (200 ms at 50% power) where acquired every 1 and 2 minutes for growth on LB and Glucose, respectively.

### Quantification and Statistical analysis

#### Analysis of time-lapse microscopy movies

Time-lapse data were processed with custom MATLAB functions developed within our group ^20,35^. Briefly, an automated pipeline segmented every image, identifying individual cells and calculating their spatial features. Cells were assigned unique identifiers and were tracked in time, allowing for the calculation of time-dependent properties including cell ages, cell sizes (areas and lengths), elongation rates and generation times. In addition, the genealogy of every cell was recorded. The fluorescence values that we report here are the sum of all pixel intensities in the area of a cell contour. As a measure for fluorescence concentration we calculated the average pixel intensity in the aread of a the cell countour (i.e. sum of all pixel intensities divided by number of pixels).

Several data filters were applied to produce a coherent data set. Firstly, data was filtered to retain only cells for which complete cell cycles, i.e. birth and division events, were observed. Therefore, cells present at the start of an experiment were eliminated, as their births were not observed. Similarly, all cells with an incomplete cell cycle at the end of the experiment are removed. Also, any cells that display a length decrease within the first 10 % of their cell cycle are flagged as segmentation errors. These cells along with their sister cell are excluded from further analysis. Lastly, we observed some filamentation in the experiment with strain QC101 growing on LB. For this experiment we applied several additional filters to remove filamenting cells and their progeny, these included: cells with a division length > 2.5 × their birth length, cells with an interdivision time > 2 × the population average, cells with an interdivision time of < 10 min, cells with a birth length > 10 *μ*m (∼ 3 × population average). In total, these criteria resulted in 594 (filamenting cells and their progeny) out of 6217 cells being excluded. See Tables S1-S3 for a summary of total cell numbers and average characteristics for each experiment.

#### Calculation of population specific growth rate

The specific growth rate of the population is calculated from the slope of a fitted linear function to the sum of the logarithm-transformed (*Ln*) lengths of all single cells.

#### Binning by normalized age and calculation of specific elongation rate

The normalized age of a cell is is calculated by dividing the absolute age (min) by the interdivision time (min). Therefore, at birth the normalized age is 0 and at division it is 1. To calculate the specific elongation rate of a single cell as a function of its normalized age, we used a piecewise approach by binning the normalized time series into 10 age bins of width 0.1 each. Next, the *Ln*-difference of the first and last data point of each age bin was taken and divided by ^1^ × interdivision time to calculate the specific elongation rate; this yielded 10 sER values per cell cycle for every single cell.

#### Binning by birth length

For the analysis of birth size-dependent cell cycle dynamics, cells were binned into classes depending on their length at birth. For each experiment, the birth length of singles cells is rescaled by division with the average birth length of all cells. Next, cells are binned into bins with a relative width of 0.075. Only bins with at least 100 individual cells are retained for further analysis. See Tables S1-S3 for details on the absolute and relative size ranges the bins, and cell numbers per bin, for each experiment.

### Growth model

Simulations for the growth model were done using Mathematica (Wolfram research). Model details are provided in the Supplementary information.

