## Supplemental information for "Growth consequences of the inhomogeneous organization of the bacterial cytoplasm"

April 17, 2023

This supplemental document contains two parts: the supplemental information concerning *i*) the experimental data and *ii*) the theoretical model. The figures referred to in this supplemental text that can be found in the main text are indicated without a prefix. The figures that can only found as supplemental figures are labelled with prefix 'S'.

### Contents

**I Data - Supplemental Information S3**

**S1 Overview of birth-size-class size ranges and cell numbers, and population-average features per experiment S11**

**II Model - Supplemental Information S14**

**S2 Preliminaries S14**

S2.1 Cell volume is approximated by the total volume of all proteins in the cell . . . . . . S14
S2.2 The specific volumetric growth rate ($\mu_V$) of single cells . . . . . . . . . . . . . . . . . . . S14
S2.3 We approximate the shape of a cell as two half-spheres (its caps) and a cylinder (mid-cell) region . . . . . . . . . . . . . . . . . . . . . . . . . . . . . . . . . . . . . . . . . . S14
S2.4 The cell length added from birth to division $\Delta L$ . . . . . . . . . . . . . . . . . . . . . . S15
S2.5 Concentration of localised proteins can differ between mother and daughter cells upon uneven division . . . . . . . . . . . . . . . . . . . . . . . . . . . . . . . . . . . . . . S15
S2.5.1 The concentration of ribosomes in single cells before and after division when they localise only in the poles (approximation) . . . . . . . . . . . . . . . . . . S16
S2.5.2 The concentration of ribosomes in single cells before and after division when they localise outside of the nucleoid (realistic model) . . . . . . . . . . . . . . . S17
S2.5.3 Estimation of cytosolic fraction occupied with ribosomes from our experimental data . . . . . . . . . . . . . . . . . . . . . . . . . . . . . . . . . . . . . . . . S19
S2.5.4 Homogenous spread of small proteins (incl. metabolic proteins) inside cells . S21

S2.5.5 The concentration of ribosomes in single cells before and after division, with volume fraction $\gamma$ filled with ribosomes . . . . . . . . . . . . . . . . . . . . . . . S21
S2.5.6 Small daughter cells grow faster than large daughter cells after birth . . . . . S22
S2.6 A deterministic model to describe the dynamics of birth-size-dependent, single-cell growth rates from birth to division . . . . . . . . . . . . . . . . . . . . . . . . . . . . . . . . S22
S2.6.1 Metabolism and protein expression model . . . . . . . . . . . . . . . . . . . . . . S22
S2.6.2 The growth-rate disturbance at cell birth is compensated for during the cell cycle by the regulatory action of ppGpp . . . . . . . . . . . . . . . . . . . . . . . S23
S2.6.3 Model design requirements . . . . . . . . . . . . . . . . . . . . . . . . . . . . . . . . . S24
S2.6.4 Including two different types of length growth makes the model comparable to the experimental data . . . . . . . . . . . . . . . . . . . . . . . . . . . . . . . . . S25
S2.6.5 Timing and initiation of two growth modes (cylindrical and midcell caps) . . S29
S2.7 Simulating thousands of cell generations to capture the non-steady state mother effect S30

### Part I

### Data - Supplemental Information

#### Confirmation of the balanced growth range

During steady state balanced growth, the specific growth rate and other properties of the population should remain fixed for a duration longer than several generation times[16]. For each experiment, we identified the period during which the population growth rate remains fixed, by plotting the natural logarithm of the sum of lengths of all cells as function of time. We then used a sliding window with a size equal to $\frac{1}{2}$ of the average interdivision time of individual cells to calculate the sliding $R^2$ value of a linear fit to the population length profile. We then identified the longest interval with an $R^2$ value bigger than 0.999 (fig. S1) to delineate the time range during which the population exhibits stable exponential growth. All subsequent analysis were then performed using data from this range. Additionally, we confirmed balanced growth during this interval by evaluating characteristics which has previously been confirmed to reflect the balanced growth status[21] of a cell population. These included the age distribution (fig. S1), which should match the theoretical distribution formulated by Painter and Marr[16], in addition to time invariant distributions of birth and division sizes (fig. S1).

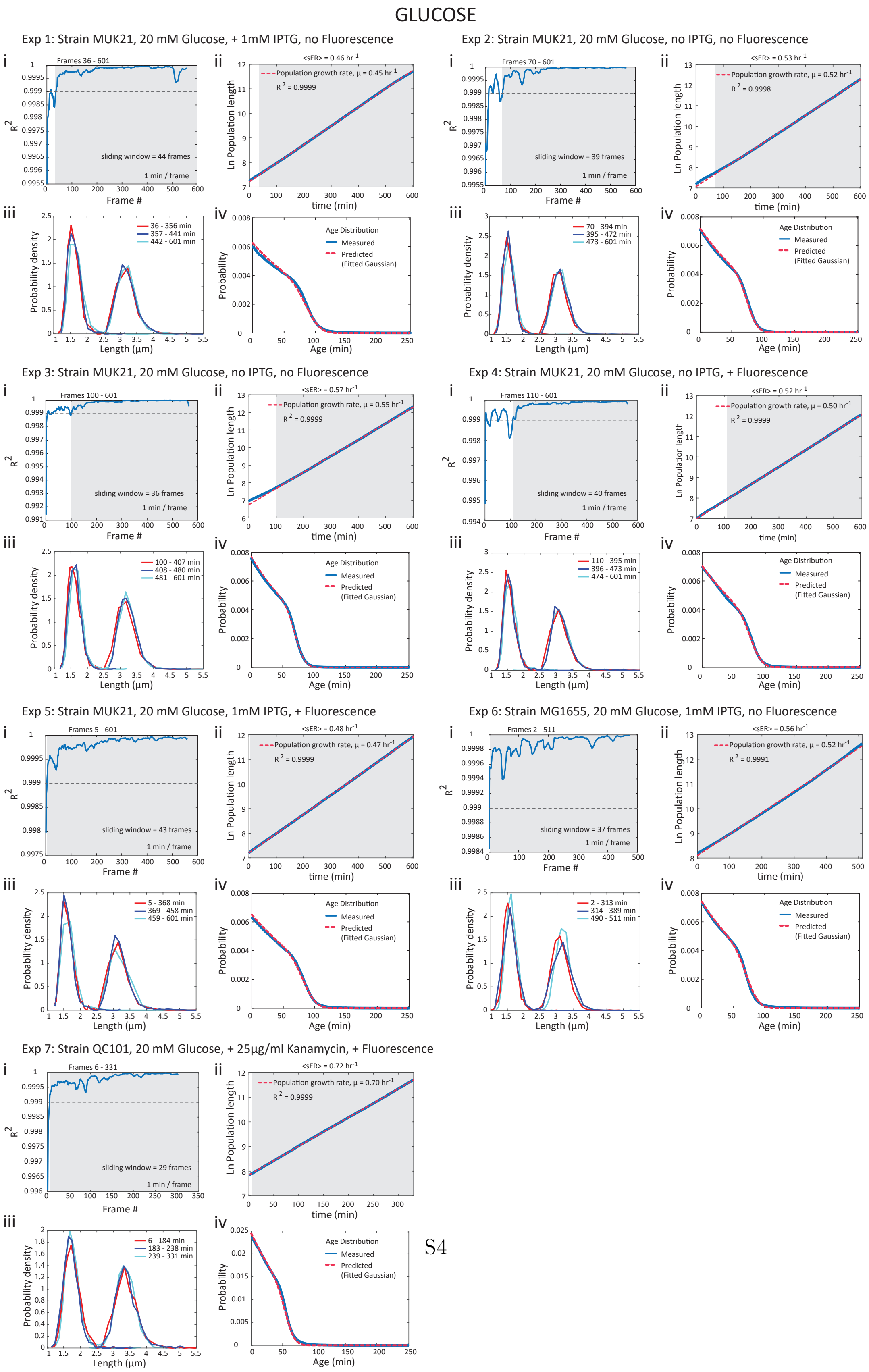


Figure S1: Figure legend on next page.

Figure S1: **Cells exhibit balanced growth on the population level.** Plots for each individual experiment are grouped (i-iv). (i) Using a sliding window of size equal to $\frac{1}{2}$ of the average interdivision time, the $R^2$ value of linear fits to the $Ln$-transformed total length profile of the population is calculated to identify the data range that displays stable exponential growth. The grey area indicates the longest continuous range with $R^2$ values above 0.999 (cutoff used). We use only data from this stable exponential growth range (analysis range) in all subsequent analyses. (ii) $Ln$-transformed total-population-length profile (blue line), with a linear fit (dashed red line) to data from the stable growth regime (grey area) indicating the specific growth rate of the population. We note that the specific growth rate of the population is in excellent agreement with the single cells $\langle sER \rangle$ (also see Table S1). (iii) Distributions of birth and division length are stable across several generations, indicating balanced population growth. Each distribution represents a third of all cells per condition from the analysis range (grey area in i and ii). (iv) The measured age distribution (blue line), data from analysis range) are compared to a theoretical distribution (dashed red line) calculated using the relation between generation time and growth rate for populations at balanced growth, as formulated by Painter and Marr[16]

### Birth-size dependent growth rate deviations of individual experiments

We performed multiple experiments, to quantify the birth-size-dependent specific elongation rate (sER) patterns under different imaging conditions (with or without fluorescence excitation), with different strains, and with different growth media (fig. S2; see Tables S1-S3 for details). We note that the growth patterns for all glucose experiments are very similar, with a clear birth size-dependence. On glucose, small differences in the relative sER deviations at birth are seen between different strains (Fig. S2, panels a-g), but overall the growth behaviour is highly reproducible. Lactose growth (fig. S2, panels h-j) shows patterns similar to glucose, while growth on LB (fig. S2, panels k and l) shows a distinctly different pattern, with no birth-size dependence.

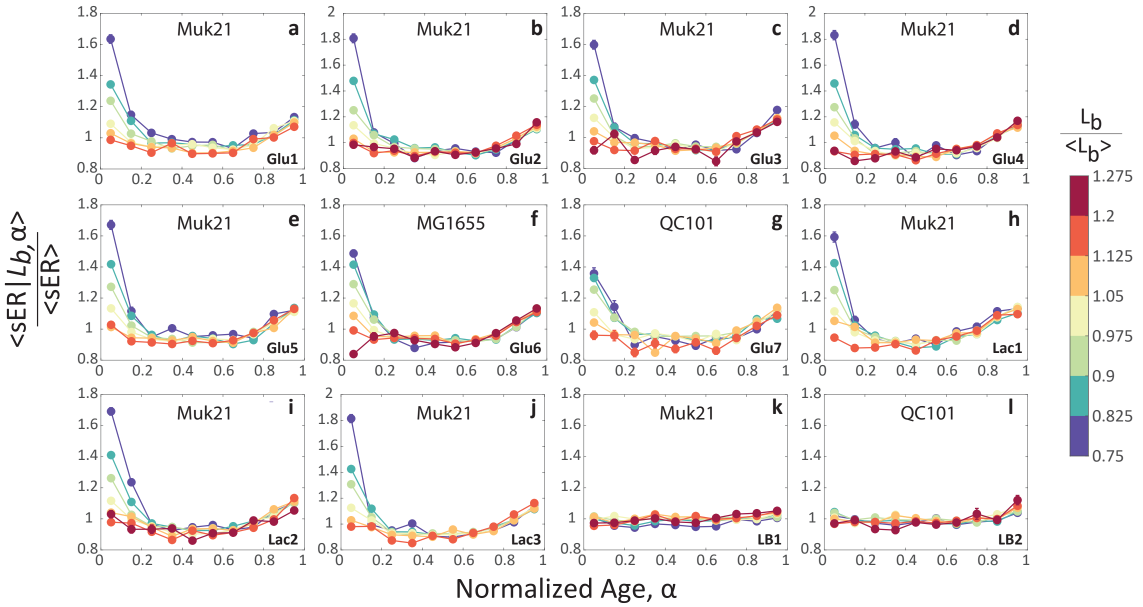


Figure S2: **Specific elongation rate profiles.** Shown are the birth length binned specific elongation rate (<sER> $|L_b$,$\alpha$) profiles for individual Glucose (a-g), Lactose (h-j) and LB (k-l) experiments, as function of normalized age ($\alpha$). Where $\alpha$=0 implies birth and 1 division. Strain names are shown. See Tables S2 - S3 for additional details.

#### Deviation and compensation of cell length

Cell length deviations are largest at birth for all conditions tested, and decrease as a function of the cell cycle (fig. S3).

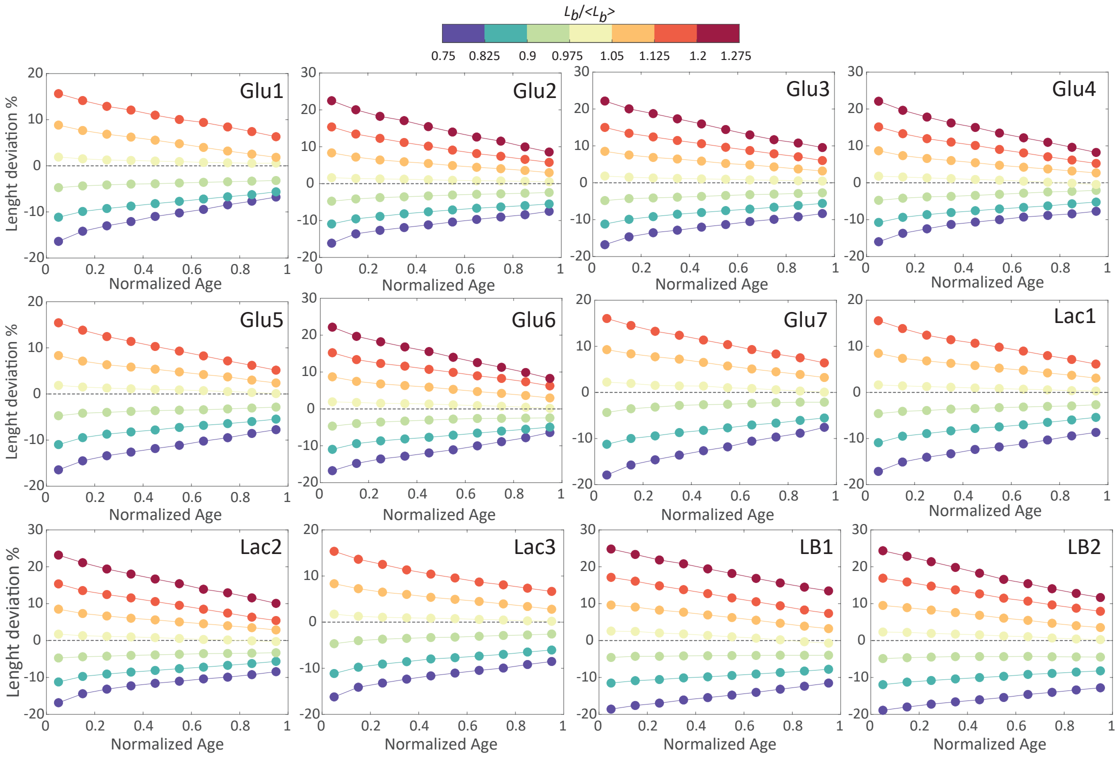


Figure S3: **Deviation and compensation of cell length.** Shown are length deviations (from the population average length) for each birth size bin, as a function of the cell cycle. For each plot, the corresponding experiment is indicated (see Tables S1-S3 for details).

#### Deviation, compensation and homeostasis of ribosomes and homogenously dispersed protein (GFP)

For both ribosomes and GFP, the total fluorescence signal at birth scales with size; larger cells have a higher signal than smaller cells (fig. S4A, B and C, left panels). Size-dependent deviations at birth are large, in the order of ~ 20 – 30% (relative to the population mean) for the largest and smallest size bins, respectively. Deviations are significantly reduced at the end of the cell cycle, but compensation is incomplete and differences remain.

For the total ribosome fluorescence per area (fluorescence concentration), we observe a distinct difference between cells growing on glucose and LB (fig. S4A and B, right panels). For cells growing on glucose, we see an inverse relation between birth size and ribosome fluoresence per area; smaller cells have a higher signal than average and larger cells lower. For cells growing on LB, we observe no size-dependence of fluorescence per area at birth, or at any time during the cell cycle. For glucose, we again see compensation during the cell cycle, but to a much lesser degree than observed for the total fluorescence signal. Importantly, the ribosome fluorescence per area pattern for LB is similar to the pattern of a homogenously dispersed GFP (fig. S4C, right panels), with no size dependence of fluorescence per area.

The degree to which ribosome concentration deviations are compensated by the end of the cell cycle, is dependent on both compensation of deviations of total ribosomes and cell size. The strong coupling between ribosome activity (protein synthesis) and mass growth means that changes in ribosome content will propagate to mass growth. In fact when looking at growth on glucose, we note the same pattern of compensation for both cell size and total ribosomes. That is, when we look at the increment of length or total ribosome fluorescence added during the cell cycle, both quantities display a sizer-like adder behaviour[23] (fig. S5A). This coupling has important consequences for the degree to which compensation of concentration deviations can be achieved during the cell cycle. For example, for smaller cells to achieve (near) full compensation of the birth deviation, the amount of ribosomes added during the cell cycle, should significantly exceed the amount of cell size added. In contrast, for larger cells, the amount of cell size added should be significantly larger than the amount of ribosomes added to achieve (near) full compensation. The concentration data shows that compensation is very modest within one cell cycle (fig. S4A, left panel), compared to the total fluorescence signal (fig. S4A, right panel). This can be understood in light of the strong correlation between cell size and total ribosome fluorescence correction, which implies that compensation of ribosome concentration deviations will be much slower than compensation of deviations in size and total ribosome content.

Lastly, the population average total fluorescence signal, for both ribosomes and GFP, doubles during the cell cycle (fig. S4D and E, left panels), while the signal per area (concentration) (fig. S4D and E, right panels), remains approximately constant.

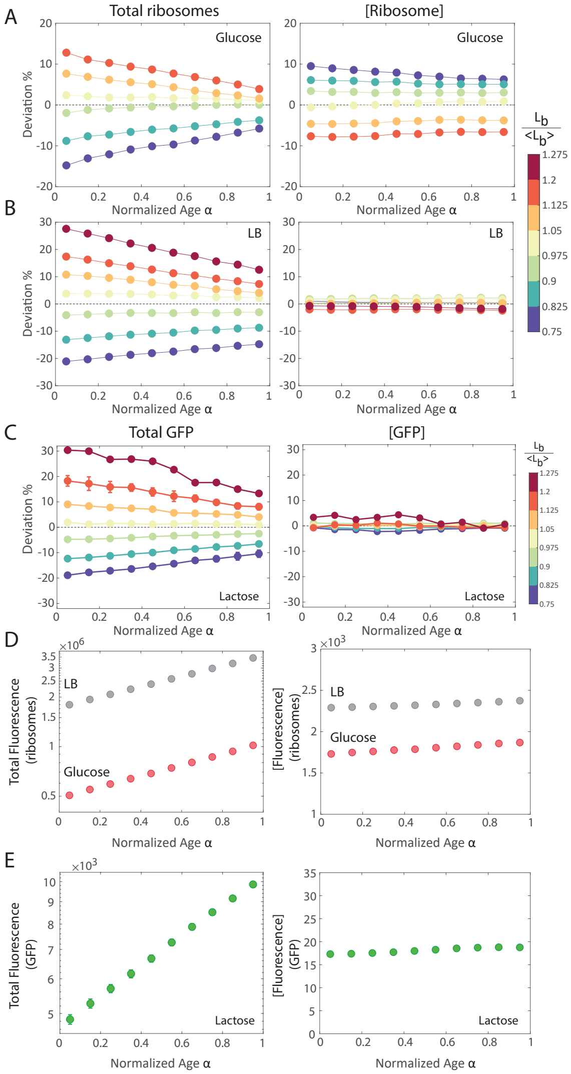


Figure S4: **Deviation, compensation and homeostasis of fluorescence.** (A) Birth size-dependent patterns of total ribosome fluorescence (left panel) and ribosome fluorescence per area (concentration) (right panel), for cells growing on glucose. (B) Birth size-dependent patterns of total ribosome fluorescence (left panel) and ribosome fluorescence per area (concentration) (right panel), for cells growing on LB. (C) Birth size-dependent patterns of total GFP fluorescence (left panel) and GFP fluorescence per area (concentration) (right panel), for cells growing on lactose. (D) Population averaged total ribosome fluorescence (left panel) and ribosome fluorescence per area (concentration) (right panel). (E) Population averaged total GFP fluorescence (left panel) and GFP fluorescence per area (concentration) (left panel).

**Comparison of cell-size control mechanisms and total cellular ribosome compensation** We note that "ribosome-growth", i.e. the amount of ribosomes added during the cell cycle, follows the same pattern as length-growth for both Glucose and LB conditions. For cells growing on glucose, this pattern corresponds to a sizer-like-adder mechanism of growth, where smaller-than-average cells grow by a slightly larger increment than the average cell, while the opposite is observed for larger-than-average cells. This pattern is similar to that reported by Wallden et al.[23] for slow growing cells. For cells growing on LB, the growth pattern corresponds to an adder-mechanism, for both length and absolute ribosome content. Again this pattern is similar to that reported by Wallden et al. for large and very fast growing cells.

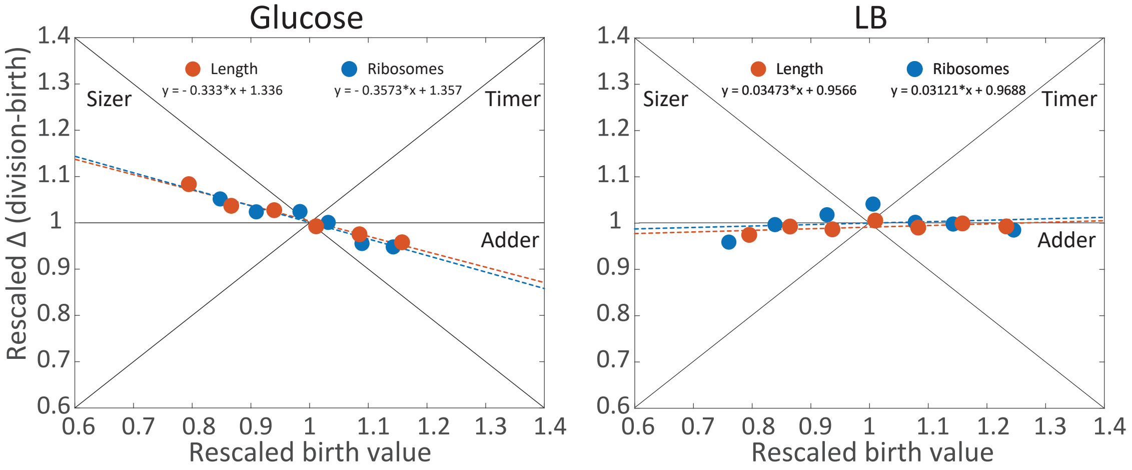


Figure S5: **Sizer, Adder, Timer relation - length and ribosomes.** Delta Ribo (Fluo) is calculated using the first and last frames containing a fluorescence value. Either can be 1 frame off. Delta length is calculated from first and last frames.

#### S1 Overview of birth-size-class size ranges and cell numbers, and population-average features per experiment

Table S1: **Summary data for Glucose experiments.** Shown are length bin class size ranges, absolute and relative, and number of unique cells per size class. Population averages for birth and division length, $\Delta$length, cell width, interdivision time, specific elongation rate and specific population growth rate are also shown.

**Experiment Glu1: Strain MUK21, 20 mM Glucose, + 1mM IPTG, no Fluorescence**

| Birth size bins | | | | | | | | |
|---|---|---|---|---|---|---|---|---|
| Bin Start ($\mu$m) | Bin End ($\mu$m) | Bin Average ($\mu$m) | Relative Start | Relative End | Relative Average | N | Population Averages | |
| 1.2281 | 1.3509 | 1.3122 | 0.75 | 0.825 | 0.8014 | 112 | <Birth Length> ($\mu$m) | 1.6374 |
| 1.35091 | 1.4737 | 1.4212 | 0.825 | 0.9 | 0.8680 | 551 | <Division Length> ($\mu$m) | 3.2050 |
| 1.47371 | 1.5965 | 1.5352 | 0.9 | 0.975 | 0.9376 | 820 | <$\Delta$Length> ($\mu$m) | 1.5676 |
| 1.59651 | 1.7193 | 1.6547 | 0.975 | 1.05 | 1.0105 | 696 | <Width> ($\mu$m) | 0.6567 |
| 1.71931 | 1.8421 | 1.7771 | 1.05 | 1.125 | 1.0853 | 537 | <Interdivision time> (min) | 88.8710 |
| 1.84211 | 1.9649 | 1.8958 | 1.125 | 1.2 | 1.1578 | 209 | <Specific elongation rate> (hr$^{-1}$) | 0.4640 |
| | | | | | | 2925 | Specific Population growth rate (hr$^{-1}$) | 0.4536 |

**Experiment Glu2: Strain MUK21, 20 mM Glucose, no IPTG, no Fluorescence**

| Birth size bins | | | | | | | | |
|---|---|---|---|---|---|---|---|---|
| Bin Start ($\mu$m) | Bin End ($\mu$m) | Bin Average ($\mu$m) | Relative Start | Relative End | Relative Average | N | Population Averages | |
| 1.1963 | 1.3159 | 1.2762 | 0.75 | 0.825 | 0.8001 | 146 | <Birth Length> ($\mu$m) | 1.5951 |
| 1.31591 | 1.4356 | 1.3846 | 0.825 | 0.9 | 0.8680 | 879 | <Division Length> ($\mu$m) | 3.1391 |
| 1.43561 | 1.5552 | 1.4988 | 0.9 | 0.975 | 0.9396 | 1524 | <$\Delta$Length> ($\mu$m) | 1.5440 |
| 1.55521 | 1.6748 | 1.6110 | 0.975 | 1.05 | 1.0100 | 1421 | <Width> ($\mu$m) | 0.6850 |
| 1.67481 | 1.7945 | 1.7282 | 1.05 | 1.125 | 1.0835 | 934 | <Interdivision time> (min) | 77.2001 |
| 1.79451 | 1.9141 | 1.8471 | 1.125 | 1.2 | 1.1580 | 399 | <Specific elongation rate> (hr$^{-1}$) | 0.5339 |
| 1.91411 | 2.0337 | 1.9640 | 1.2 | 1.275 | 1.2313 | 140 | Specific Population growth rate (hr$^{-1}$) | 0.5196 |
| | | | | | | 5443 | | |

**Experiment Glu3: Strain MUK21, 20 mM Glucose, no IPTG, no Fluorescence**

| Birth size bins | | | | | | | | |
|---|---|---|---|---|---|---|---|---|
| Bin Start ($\mu$m) | Bin End ($\mu$m) | Bin Average ($\mu$m) | Relative Start | Relative End | Relative Average | N | Population Averages | |
| 1.2243 | 1.3468 | 1.3098 | 0.75 | 0.825 | 0.8023 | 185 | <Birth Length> ($\mu$m) | 1.6325 |
| 1.34681 | 1.4692 | 1.4171 | 0.825 | 0.9 | 0.8681 | 914 | <Division Length> ($\mu$m) | 3.2065 |
| 1.46921 | 1.5916 | 1.5328 | 0.9 | 0.975 | 0.9390 | 1407 | <$\Delta$Length> ($\mu$m) | 1.5740 |
| 1.59161 | 1.7141 | 1.6516 | 0.975 | 1.05 | 1.0117 | 1416 | <Width> ($\mu$m) | 0.6690 |
| 1.71411 | 1.8365 | 1.7706 | 1.05 | 1.125 | 1.0846 | 927 | <Interdivision time> (min) | 72.1765 |
| 1.83651 | 1.9590 | 1.8855 | 1.125 | 1.2 | 1.1550 | 477 | <Specific elongation rate> (hr$^{-1}$) | 0.5704 |
| 1.95901 | 2.0814 | 2.0080 | 1.2 | 1.275 | 1.2300 | 163 | Specific Population growth rate (hr$^{-1}$) | 0.5534 |
| | | | | | | 5489 | | |

**Experiment Glu4: Strain MUK21, 20 mM Glucose, no IPTG, + Fluorescence**

| Birth size bins | | | | | | | | |
|---|---|---|---|---|---|---|---|---|
| Bin Start ($\mu$m) | Bin End ($\mu$m) | Bin Average ($\mu$m) | Relative Start | Relative End | Relative Average | N | Population Averages | |
| 1.1904 | 1.3094 | 1.2736 | 0.75 | 0.825 | 0.8024 | 125 | <Birth Length> ($\mu$m) | 1.5875 |
| 1.30941 | 1.4285 | 1.3805 | 0.825 | 0.9 | 0.8698 | 751 | <Division Length> ($\mu$m) | 3.1233 |
| 1.42851 | 1.5475 | 1.4885 | 0.9 | 0.975 | 0.9378 | 1195 | <$\Delta$Length> ($\mu$m) | 1.5361 |
| 1.54751 | 1.6665 | 1.6050 | 0.975 | 1.05 | 1.0112 | 1079 | <Width> ($\mu$m) | 0.6937 |
| 1.66651 | 1.7856 | 1.7214 | 1.05 | 1.125 | 1.0845 | 638 | <Interdivision time> (min) | 79.2804 |
| 1.78561 | 1.9046 | 1.8368 | 1.125 | 1.2 | 1.1573 | 334 | <Specific elongation rate> (hr$^{-1}$) | 0.5208 |
| 1.90461 | 2.0237 | 1.9537 | 1.2 | 1.275 | 1.2309 | 130 | Specific Population growth rate (hr$^{-1}$) | 0.5039 |
| | | | | | | 4252 | | |

**Experiment Glu5: Strain MUK21, 20 mM Glucose, + 1mM IPTG, + Fluorescence**

| Birth size bins | | | | | | | | |
|---|---|---|---|---|---|---|---|---|
| Bin Start ($\mu$m) | Bin End ($\mu$m) | Bin Average ($\mu$m) | Relative Start | Relative End | Relative Average | N | Population Averages | |
| 1.2206 | 1.3427 | 1.3057 | 0.75 | 0.825 | 0.8023 | 149 | <Birth Length> ($\mu$m) | 1.6275 |
| 1.34271 | 1.4648 | 1.4127 | 0.825 | 0.9 | 0.8680 | 617 | <Division Length> ($\mu$m) | 3.1900 |
| 1.46481 | 1.5868 | 1.5266 | 0.9 | 0.975 | 0.9380 | 1052 | <$\Delta$Length> ($\mu$m) | 1.5625 |
| 1.58681 | 1.7089 | 1.6446 | 0.975 | 1.05 | 1.0105 | 891 | <Width> ($\mu$m) | 0.6695 |
| 1.70891 | 1.8309 | 1.7601 | 1.05 | 1.125 | 1.0815 | 613 | <Interdivision time> (min) | 85.2068 |
| 1.83091 | 1.8309 | 1.8837 | 1.125 | 1.2 | 1.1574 | 286 | <Specific elongation rate> (hr$^{-1}$) | 0.4837 |
| | | | | | | 3608 | Specific Population growth rate (hr$^{-1}$) | 0.4702 |

**Experiment Glu6: Strain MG1655, 20 mM Glucose, no IPTG, no Fluorescence**

| Birth size bins | | | | | | | | |
|---|---|---|---|---|---|---|---|---|
| Bin Start ($\mu$m) | Bin End ($\mu$m) | Bin Average ($\mu$m) | Relative Start | Relative End | Relative Average | N | Population Averages | |
| 1.1980 | 1.3178 | 1.2794 | 0.75 | 0.825 | 0.8010 | 161 | <Birth Length> ($\mu$m) | 1.5974 |
| 1.31781 | 1.4376 | 1.3879 | 0.825 | 0.9 | 0.8689 | 665 | <Division Length> ($\mu$m) | 3.1624 |
| 1.43761 | 1.5574 | 1.4996 | 0.9 | 0.975 | 0.9388 | 1143 | <$\Delta$Length> ($\mu$m) | 1.5651 |
| 1.55741 | 1.6772 | 1.6151 | 0.975 | 1.05 | 1.0111 | 966 | <Width> ($\mu$m) | 0.6974 |
| 1.67721 | 1.7970 | 1.7311 | 1.05 | 1.125 | 1.0837 | 674 | <Interdivision time> (min) | 74.7321 |
| 1.79701 | 1.9168 | 1.8451 | 1.125 | 1.2 | 1.1551 | 319 | <Specific elongation rate> (hr$^{-1}$) | 0.5606 |
| 1.91681 | 2.0366 | 1.9707 | 1.2 | 1.275 | 1.2337 | 109 | Specific Population growth rate (hr$^{-1}$) | 0.5212 |
| | | | | | | 7462 | | |

**Experiment Glu7: Strain QC101, 20 mM Glucose, + 25$\mu$g/ml Kanamycin, + Fluorescence**

| Birth size bins | | | | | | | | |
|---|---|---|---|---|---|---|---|---|
| Bin Start ($\mu$m) | Bin End ($\mu$m) | Bin Average ($\mu$m) | Relative Start | Relative End | Relative Average | N | Population Averages | |
| 1.3197 | 1.4517 | 1.3959 | 0.75 | 0.825 | 0.7933 | 154 | <Birth Length> ($\mu$m) | 1.7596 |
| 1.45171 | 1.5836 | 1.5231 | 0.825 | 0.9 | 0.8656 | 450 | <Division Length> ($\mu$m) | 3.3915 |
| 1.58361 | 1.7156 | 1.6523 | 0.9 | 0.975 | 0.9390 | 669 | <$\Delta$Length> ($\mu$m) | 1.6319 |
| 1.71561 | 1.8476 | 1.7781 | 0.975 | 1.05 | 1.0105 | 623 | <Width> ($\mu$m) | 0.7043 |
| 1.84761 | 1.9795 | 1.9082 | 1.05 | 1.125 | 1.0845 | 430 | <Interdivision time> (min) | 57.6394 |
| 1.97951 | 2.1115 | 2.0362 | 1.125 | 1.2 | 1.1572 | 243 | <Specific elongation rate> (hr$^{-1}$) | 0.7163 |
| | | | | | | 2569 | Specific Population growth rate (hr$^{-1}$) | 0.6982 |

Table S2: **Summary data for Lactose experiments.** Shown are length bin class size ranges, absolute and relative, and number of unique cells per size class. Population averages for birth and division length, $\Delta$length, cell width, interdivision time, specific elongation rate and specific population growth rate are also shown.

**Experimental Lac1: 10 mM Lactose, 1mM IPTG, + Fluorescence**

| Birth size bins | | | | | | | | |
|---|---|---|---|---|---|---|---|---|
| Bin Start ($\mu$m) | Bin End ($\mu$m) | Bin Average ($\mu$m) | Relative Start | Relative End | Relative Average | N | Population Averages | |
| 1.3268 | 1.4594 | 1.4129 | 0.825 | 0.9 | 0.7987 | 111 | <Birth Length> ($\mu$m) | 1.7690 |
| 1.45941 | 1.5921 | 1.5367 | 0.9 | 0.975 | 0.8687 | 359 | <Division Length> ($\mu$m) | 3.4186 |
| 1.59211 | 1.7248 | 1.6641 | 0.975 | 1.05 | 0.9407 | 709 | <$\Delta$Length> ($\mu$m) | 1.6496 |
| 1.72481 | 1.8575 | 1.7871 | 1.05 | 1.125 | 1.0103 | 746 | <Width> ($\mu$m) | 0.7025 |
| 1.85751 | 1.9901 | 1.9164 | 1.125 | 1.2 | 1.0833 | 440 | <Interdivision time> (min) | 79.8585 |
| 1.99011 | 2.1228 | 2.0492 | 1.2 | 1.275 | 1.1584 | 191 | <Specific elongation rate> (hr$^{-1}$) | 0.5013 |
| | | | | | | 2556 | Specific Population growth rate (hr$^{-1}$) | 0.4994 |

**Experimental Lac2: 10 mM Lactose, no IPTG, + Fluorescence**

| Birth size bins | | | | | | | | |
|---|---|---|---|---|---|---|---|---|
| Bin Start ($\mu$m) | Bin End ($\mu$m) | Bin Average ($\mu$m) | Relative Start | Relative End | Relative Average | N | Population Averages | |
| 1.2448 | 1.3692 | 1.3276 | 0.75 | 0.825 | 0.7999 | 232 | <Birth Length> ($\mu$m) | 1.6597 |
| 1.36921 | 1.4937 | 1.4384 | 0.825 | 0.9 | 0.8667 | 776 | <Division Length> ($\mu$m) | 3.2729 |
| 1.49371 | 1.6182 | 1.5577 | 0.9 | 1.05 | 0.9385 | 1260 | <$\Delta$Length> ($\mu$m) | 1.6132 |
| 1.61821 | 1.7427 | 1.6775 | 0.975 | 1.05 | 1.0107 | 1230 | <Width> ($\mu$m) | 0.6939 |
| 1.74271 | 1.8671 | 1.7981 | 1.05 | 1.125 | 1.0834 | 845 | <Interdivision time> (min) | 86.8290 |
| 1.86711 | 1.9916 | 1.9194 | 1.125 | 1.2 | 1.1565 | 357 | <Specific elongation rate> (hr$^{-1}$) | 0.4777 |
| 1.99161 | 2.1161 | 2.0483 | 1.2 | 1.275 | 1.2341 | 164 | Specific Population growth rate (hr$^{-1}$) | 0.4799 |
| | | | | | | 4864 | | |

**Experimental Lac3: 10 mM Lactose, 1mM IPTG, + Fluorescence**

| Birth size bins | | | | | | | | |
|---|---|---|---|---|---|---|---|---|
| Bin Start ($\mu$m) | Bin End ($\mu$m) | Bin Average ($\mu$m) | Relative Start | Relative End | Relative Average | N | Population Averages | |
| 1.2118 | 1.3330 | 1.2726 | 0.75 | 0.825 | 0.7875 | 166 | <Birth Length> ($\mu$m) | 1.6158 |
| 1.33301 | 1.4542 | 1.3938 | 0.825 | 0.9 | 0.8625 | 705 | <Division Length> ($\mu$m) | 3.1878 |
| 1.45421 | 1.5754 | 1.5150 | 0.9 | 1.05 | 0.975 | 1043 | <$\Delta$Length> ($\mu$m) | 1.5720 |
| 1.57541 | 1.6966 | 1.6362 | 0.975 | 1.05 | 1.0125 | 1097 | <Width> ($\mu$m) | 0.7181 |
| 1.69661 | 1.8178 | 1.7574 | 1.05 | 1.125 | 1.0875 | 696 | <Interdivision time> (min) | 83.4236 |
| 1.81781 | 1.9390 | 1.8786 | 1.125 | 1.2 | 1.1625 | 364 | <Specific elongation rate> (hr$^{-1}$) | 0.4967 |
| | | | | | | 4186 | Specific Population growth rate (hr$^{-1}$) | 0.4703 |

Table S3: **Summary data for LB experiments.** Shown are length bin class size ranges, absolute and relative, and number of unique cells per size class. Population averages for birth and division length, $\Delta$length, cell width, interdivision time, specific elongation rate and specific population growth rate are also shown.

**Experimental LB1: Strain MG1655, LB, no IPTG, no Fluorescence**

| Birth size bins | | | | | | | | |
|---|---|---|---|---|---|---|---|---|
| Bin Start ($\mu$m) | Bin End ($\mu$m) | Bin Average ($\mu$m) | Relative Start | Relative End | Relative Average | N | Population Averages | |
| 2.5264 | 2.7791 | 2.6813 | 0.75 | 0.825 | 0.7960 | 405 | <Birth Length> ($\mu$m) | 3.3686 |
| 2.77911 | 3.0317 | 2.9236 | 0.825 | 0.9 | 0.8679 | 991 | <Division Length> ($\mu$m) | 6.4608 |
| 3.03171 | 3.2844 | 3.1596 | 0.9 | 1.05 | 0.9380 | 1434 | <$\Delta$Length> ($\mu$m) | 3.0922 |
| 3.28441 | 3.5370 | 3.4051 | 0.975 | 1.05 | 1.0109 | 1321 | <Width> ($\mu$m) | 0.9835 |
| 3.53701 | 3.7896 | 3.6448 | 1.05 | 1.125 | 1.0820 | 841 | <Interdivision time> (min) | 23.0435 |
| 3.78961 | 4.0423 | 3.9053 | 1.125 | 1.2 | 1.1593 | 406 | <Specific elongation rate> (hr$^{-1}$) | 1.7153 |
| 4.04231 | 4.0423 | 4.1581 | 1.2 | 1.275 | 1.2344 | 191 | Specific Population growth rate (hr$^{-1}$) | 1.7198 |
| | | | | | | 5589 | | |

**Experimental LB2: Strain QC101, LB, + 25$\mu$g/ml Kanamycin, + Fluorescence**

| Birth size bins | | | | | | | | |
|---|---|---|---|---|---|---|---|---|
| Bin Start ($\mu$m) | Bin End ($\mu$m) | Bin Average ($\mu$m) | Relative Start | Relative End | Relative Average | N | Population Averages | |
| 2.4833 | 2.7316 | 2.6330 | 0.75 | 0.825 | 0.7952 | 232 | <Birth Length> ($\mu$m) | 3.3111 |
| 2.73161 | 2.9800 | 2.8632 | 0.825 | 0.9 | 0.8647 | 776 | <Division Length> ($\mu$m) | 6.3495 |
| 2.98001 | 3.2283 | 3.1023 | 0.9 | 1.05 | 0.9370 | 1260 | <$\Delta$Length> ($\mu$m) | 3.0384 |
| 3.22831 | 3.4766 | 3.3440 | 0.975 | 1.05 | 1.0100 | 1230 | <Width> ($\mu$m) | 0.9972 |
| 3.47661 | 3.7250 | 3.5864 | 1.05 | 1.125 | 1.0832 | 845 | <Interdivision time> (min) | 25.7386 |
| 3.72501 | 3.9733 | 3.8358 | 1.125 | 1.2 | 1.1585 | 357 | <Specific elongation rate> (hr$^{-1}$) | 1.5389 |
| 3.97331 | 4.2216 | 4.0834 | 1.2 | 1.275 | 1.2333 | 164 | Specific Population growth rate (hr$^{-1}$) | 1.5193 |
| | | | | | | 4864 | | |

### Part II

### Model - Supplemental Information

#### S2 Preliminaries

##### S2.1 Cell volume is approximated by the total volume of all proteins in the cell

We assume that the volume of a cell, $V$, equals the sum of the volumes of its proteins,

$$V = \sum_{i \in \text{proteins}} \hat{v}_i n_i \tag{1}$$

with $\hat{v}_i$ as the volume of protein $i$ and $n_i$ as its copy number in the cell. The volume of a protein can be enlarged with a cytosolic shell to accommodate the composition of the cytosol that are not proteins.

##### S2.2 The specific volumetric growth rate ($\mu_V$) of single cells

We define the specific growth rate of the volume of a cell, $\mu_V$, in terms of the protein synthesis rate, which is a consensus modelling assumption in the field[19],

$$\mu_V = \frac{1}{V}\dot{V} = \sum_i \hat{v}_i \frac{1}{V}\dot{n}_i = \sum_i \hat{v}_i \frac{1}{V}\alpha_i k_r f_r n_r = \sum_i \hat{v}_i \alpha_i k_r f_r c_r, \tag{2}$$

with $\alpha_i k_r f_r c_r$ as the synthesis rate of protein $i$ by the ribosome. The fraction of ribosomes translating protein $i$ equals $\alpha_i$; $k_r$ is the translation rate constant (in proteins per time); $f_r$ is a kinetic function, describing the saturation of ribosomes with its substrates; $n_r$ represents the number of ribosomes per cell; $c_r = n_r/V$ equals the concentration of ribosomes in the cell, and $c_{tot} = n/V$ equals the concentration of total protein concentration in the cell.

Assuming all modelled proteins have the same effective volume, $\hat{v}$, we obtain,

$$\mu_V = \hat{v} k_r f_r c_r = k_r f_r \frac{n_r}{V/\hat{v}} = k_r f_r \frac{n_r}{n} = k_r f_r \frac{c_r}{c_{tot}}, \tag{3}$$

##### S2.3 We approximate the shape of a cell as two half-spheres (its caps) and a cylinder (mid-cell) region

We approximate the shape of a cell (one that is not yet forming its midcell caps, prior to division) as two half-spheres (its caps) and a cylinder (mid-cell) region such that the volume of the cell can be expressed in terms its length $L$ and its radius $r$,

$$V(L) = V_{midcell} + V_{poles} = \pi r^2 (L - 2r) + \frac{4}{3}\pi r^3, \tag{4}$$

the length of the midcell region equals $L_{cyl} = L - 2r$ (Fig. S9). In our experiments we characterise the growth of the cell in terms of its length growth. At a constant radius (cylindrical growth; prior to cap formation midcell for cell division), the specific growth rate in terms of the volume $\mu_V$ and in terms of the length $\mu_V$ are related as,

$$\mu_V = \frac{1}{V}\frac{dV}{dt} = \frac{L}{V}\frac{\partial V}{\partial L}\frac{1}{L}\frac{dL}{dt} = \frac{L}{V}\frac{\partial V}{\partial L}\mu_L \tag{5}$$

Below we continue with this equation.

#### S2.4 The cell length added from birth to division $\Delta L$

According to the adder principle, the added length during one cell cycle equals,

$$\Delta L = \frac{\langle L_M \rangle}{2},$$

while the length added according to the sizer principle equals,

$$\Delta L = \langle L_M \rangle - L_B,$$

with $\langle L_M \rangle$ as the mean cell length of a dividing (mother) cell. The cells in our data show mixed adder and sizer behaviour (see also Wallden et al.[22] and Nordholt et al.[15]). We observed that $\Delta L$ decreases with $L_B$ as in the sizer theory, but less strongly so and we observe that $L_M$ increases with $L_B$, in line with the adder theory. The $\Delta L$ of our model therefore combines the two principles, with a constant added length plus a small deviation depending on the daughter's birth size. For a single daughter cell, the length added is determined by its birth length by,

$$\Delta L = \frac{\langle LM \rangle}{2} + \frac{\frac{\langle LM \rangle}{2} - L_B}{4}. \tag{6}$$

When $L_B > \frac{L_M}{2}$ this results in $\Delta L < \frac{\langle LM \rangle}{2}$ and vice versa, while still maintaining an increase in $\Delta L$ for increasing birth lengths. This way the model shows the sizer-like adder behaviour observed in the experimental data (see S5).

#### S2.5 Concentration of localised proteins can differ between mother and daughter cells upon uneven division

Large protein complexes, such as fully-assembled ribosomes and (misfolded) protein aggregates, preferentially localise outside of the mid-cell positioned nucleoid region of the cell due to an entropic force (volume-exclusion)[1,2,13,24]. On average, the mother cell will split exactly mid-cell, giving rise to two equally-sized daughters. However, as with most cellular processes, division is imperfect and noisy. This can lead to two sibling, daughter cells, each with a concentration of a localised protein that differs between them and their mother cell, while the concentration of homogeneously spread proteins is unaffected by uneven division. To see this, partition a cell into evenly-sized, volume elements and mark those that contain a specific protein green and those without red. In the case of localised proteins, we have green and red volume elements, while we only have green volume elements in the case of homogeneously spread proteins. In the latter case, regardless of how many volume elements a daughter cell obtains they are all green, they all have the same concentration. But this is not so when the protein is localised then daughter cells can receive different numbers of green and red volume elements depending on how the mother cell divides such that the concentration depends on their size and how the filled boxed where placed in the mother cell. In other words, since ribosomes are excluded from the nucleoid and metabolic proteins not, we can expect a concentration imbalance of those types of proteins when a mother cell divides (unevenly).

###### S2.5.1 The concentration of ribosomes in single cells before and after division when they localise only in the poles (approximation)

For intermediate sized cells, most ribosomes are present in the cell's poles (see fig. 2B, main text); since the volume surrounding the nucleoid in the midcell region is so small. The ribosome concentration in a daughter cell depends then on whether the mother cell divided evenly or not. When we assume that all the mother cells have the same ribosome concentration, $c_r^M$, (which is not the case in our experimental data) then the growth rate of the mother cell equals,

$$\mu_V^M = k_r f_r c_r^M \frac{1}{c_{tot}}. \tag{7}$$

Note that the saturation of the ribosome with substrates, denoted by $0 < f_r < 1$, depends on concentration of amino acids which localise homogeneously and is therefore unaffected by division and, in addition, the cell regulates the ribosome concentration to try and keep this saturation constant[4]. Thus, we assume also that all mother cells have the same ribosome saturation. These two assumptions ensure that all mother cells grow equally fast when they divide. Consider now that all ribosomes are in the poles and that each daughter cell gets one pole, each filled with the same number of ribosomes. A mother cell now contains four half-spheres (caps), two at its poles and two roughly midcell, in between her two segregated nucleoids.

The number of ribosomes of the mother (in her poles) equals

$$n_r^M = c_r^M V_M = c_r^M \left(2\frac{4}{3}\pi r^3 + \pi r^2 (L_M - 4r)\right).$$

The number of ribosomes in the daughter equals

$$n_r^D = c_r^D V_D = c_r^D \left(\frac{4}{3}\pi r^3 + \pi r^2 (L_D - 2r)\right).$$

Since all the ribosomes in the mother are evenly distributed over her two poles, each daughter gets the same of number ribosomes,

$$\frac{n_r^M}{2} = n_r^{D_1} = n_r^{D_2}.$$

From the last three equation we deduce that,

$$c_r^D = \frac{3L_M - 4r}{6L_D - 4r} c_r^M \tag{8}$$

The concentration of ribosomes in a daughter cell equals that of her mother only when she divided evenly. Then the volumes of the daughter cells are the same and half the volume of the mother cell,

$$\frac{V_M}{2} = V_{D_1} = V_{D_2} = \frac{2\frac{4}{3}\pi r^3 + \pi r^2(L_M - 4r)}{2} = \frac{4}{3}\pi r^3 + \pi r^2\left(\frac{L_M}{2} - 2r\right)$$

since the daughter always receive the same number of ribosomes, their ribosome concentration is the same upon equal division. Now consider,

$$V_M = 2\frac{4}{3}\pi r^3 + \pi r^2(\underbrace{L_{D_1} + L_{D_2}}_{L_M} - 4r) = \underbrace{\frac{4}{3}\pi r^3 + \pi r^2(L_{D_1} - 2r)}_{V_{D_1}} + \underbrace{\frac{4}{3}\pi r^3 + \pi r^2(L_{D_2} - 2r)}_{V_{D_2}},$$

indicating that the shortest daughter cell has the smallest volume and therefore the highest ribosome concentration when its received ribosome content is the same for the two daughters. For a protein that is homogeneously spread, the received content upon division is proportional to its volume, so two daughter cells always have concentrations that identical to each other and their mother – i.e. of abundant proteins with negligible noise, such as amino-acid tRNA's (substrates of the ribosome setting $f_r$ and metabolic proteins).

##### S2.5.2 The concentration of ribosomes in single cells before and after division when they localise outside of the nucleoid (realistic model)

In the previous section, we considered the simplifying case where the ribosomes are exclusively found in the poles of the cell. However, in reality there is also a fraction of the ribosomes located in the rest of the cell's cytoplasm, surrounding the midcell-positioned nucleoid. The volume fraction, $\alpha$, of the cell's mid-cell region (its cylinderical region) that contains ribosomes, so the extranucleoid midcell volume, will affect the magnitude of the birth-length-dependent disturbance of the ribosome (and growth rate) of daughter cells upon birth. A lower $\alpha$ fraction implies a low concentration of ribosomes in the cylindrical region, so then ribosomes are found mostly the poles.

In order to understand how the ribosome concentration in newborn cells depends both on the volume fraction $\alpha$ and the birth length, we consider the ribosomes concentration in the daughter ($c_D$) divided by the average concentration,

$$\begin{aligned}\frac{c_D}{\langle c_D \rangle} &= \frac{\frac{n_D}{V_D}}{\frac{\langle n_D \rangle}{\langle V_D \rangle}} \\ &= \frac{n_D}{\langle n_D \rangle}\frac{\langle V_D \rangle}{V_D},\end{aligned} \tag{9}$$

To obtain $n_D$, we will consider the number of ribosomes in a growing cell ($n$), which did not yet start to form caps in its midcell region by septum formation, which equals the sum of the ribosome number in the poles and the midcell region,

$$n = n_p + n_m$$

Next, we ensure that there exists no ribosome concentration gradient in the cell's volume that contains ribosomes; the ribosome concentration in the cell's poles equals that of extra-nucleoid midcell region,

$$\frac{n_p}{\frac{4}{3}\pi r^3} = \frac{n_m}{\alpha \pi r^2 (L-2r)} \quad \Rightarrow \quad n_m = \frac{\alpha \pi r^2 (L-2r)}{\frac{4}{3}\pi r^3}$$

such that we obtain

$$n = n_p + \alpha n_p \frac{\pi r^2 \left(L-2r\right)}{\frac{4}{3}\pi r^3} \quad \text{(number of ribosomes in a growing cell before septum formation)}. \tag{10}$$

Now we consider the average mother cell right before division when it has a length $\langle L_M \rangle$ and 4 caps instead of 2 as near-complete invagination has taken place. This results in a number of ribosomes $(n_M)$ that can be determined by,

$$n_M = 2n_p + \alpha n_p \frac{\pi r^2 \left(\langle L_M \rangle - 4r\right)}{\frac{4}{3}\pi r^3},$$

A mother that splits evenly leads to a daughter that has a ribosome copy number that equals,

$$\langle n_D \rangle = n_p + \alpha n_p \frac{\pi r^2 \left(\frac{\langle L_M \rangle}{2} - 2r\right)}{\frac{4}{3}\pi r^3}; \ \langle L_D \rangle = \frac{\langle L_M \rangle}{2},$$

note that this equation is identical to equation 10 as expected. $\langle L_D \rangle$ (strictly $\langle L_B \rangle$) equals the average daughter length at birth which equals half of the average mother length $\langle L_M \rangle$. Different daughter lengths $L_D$, due to a deviating mother length of $\delta_M$ and a deviating birth length of $\delta_B$ will give,

$$\begin{aligned}
\frac{n_D}{\langle n_D \rangle} &= \frac{n_p + \alpha n_p \frac{\pi r^2}{\frac{4}{3}\pi r^3}\left(\frac{\langle L_M \rangle + \delta_M}{2} + \delta_B - 2r\right)}{n_p + \alpha n_P \frac{\pi r^2 \left(\frac{\langle L_M \rangle}{2} - 2r\right)}{\frac{4}{3}\pi r^3}} \\
&= \frac{\frac{4}{3} r + \alpha \left(\frac{\langle L_M \rangle}{2} - 2r + \frac{\delta_M}{2} + \delta_B\right)}{\frac{4}{3} r + \alpha \left(\frac{\langle L_M \rangle}{2} - 2r\right)} \\
&= \frac{\frac{4}{3} r + \alpha \frac{\langle L_M \rangle}{2} \left(1 - \frac{4r}{\langle L_M \rangle} + \frac{2\left(\frac{\delta_M}{2} + \delta_B\right)}{\langle L_M \rangle}\right)}{\frac{4}{3} r + \alpha \frac{\langle L_M \rangle}{2} \left(1 - \frac{4r}{\langle L_M \rangle}\right)} \\
&= \frac{\frac{4}{3} r + \alpha \frac{\langle L_M \rangle}{2} \left(1 - \frac{4r}{\langle L_M \rangle} + \frac{\delta L_B}{\langle L_B \rangle}\right)}{\frac{4}{3} r + \alpha \frac{\langle L_M \rangle}{2} \left(1 - \frac{4r}{\langle L_M \rangle}\right)}
\end{aligned} \tag{11}$$

Remember from (9) that $\frac{c_D}{\langle c_D \rangle}$ is simply $\frac{n_D}{\langle n_D \rangle} \frac{\langle V_D \rangle}{V_D}$, which gives,

$$\frac{c_D}{\langle c_D\rangle} = \frac{\frac{4}{3}r + \alpha\frac{\langle L_M\rangle}{2}\left(1 - \frac{4r}{\langle L_M\rangle} + \frac{\delta L_B}{\langle L_B\rangle}\right)}{\frac{4}{3}r + \alpha\frac{\langle L_M\rangle}{2}\left(1 - \frac{4r}{\langle L_M\rangle}\right)} \times \frac{\frac{4}{3}\pi r^3 + \pi r^2\left(\frac{\langle L_M\rangle}{2} - 2r\right)}{\frac{4}{3}\pi r^3 + \pi r^2\left(L_D - 2r\right)}$$

$$= \frac{(4r - 3\langle L_M\rangle)(4\langle L_B\rangle r(2-3\alpha) + 3L_B\langle L_M\rangle\alpha)}{2\langle L_B\rangle(3L_D - 2r)(-3\langle L_M\rangle\alpha + 4r(3\alpha - 2))} \quad (12)$$

When $\alpha = 0$ we obtain $c_r^D = \frac{n_r^M/2}{V} = \frac{3\langle L_M\rangle - 4r}{6L_B - 4r}c_r^M$, which indeed equals equation 8 for the case that all ribosomes are exclusively in the poles. At $\alpha = 1$, the concentration of ribosomes is equal for all different birth lengths and $c_D/\langle c_D\rangle = 1$. Then, the ribosomes are distributed homogeneously and the concentration in the daughter cell does not depend on its birth length and is always equal to the concentration in the mother.

Since the growth rate (eq. 2) is proportional to the ribosome concentration, equations 11 and 12 will be used later to understand the growth rate of newborn cells as function of this size, given the size dependency of the ribosome copy number and concentration as derived in this section. Next, we will estimate the ribosome-filled volume fraction of the cell from our measurement of $n_D/\langle n_D\rangle$ as function of birth length.

###### S2.5.3 Estimation of cytosolic fraction occupied with ribosomes from our experimental data

To estimate the ribosome-filled volume fraction from our measurements of the birth length of single cells and their ribosome content at birth (whole cell fluorescence, not normalised per area), we fitted equation (11) to the experimental data associated with the first age bin (age bin: 0-0.05). Since we know all parameters entering the equation from measurements, the only unknown paramater is $\alpha$. The best data fit is obtained with $\alpha = 0.31$ (glucose dataset) and 0.66 (LB dataset).

From $\alpha$, which is the ribosome-occupied, volume fraction of the midcell, cylindrical region, we calculate the ribosome-filled volume fraction of the entire cell, we which we denote by $\gamma$. The relation between $\alpha$ and $\gamma$ is,

$$\text{volume fraction occupied by ribosomes} = \frac{\text{occupied volume poles}}{\text{total cell volume}} + \frac{\text{occpied volume cylinder}}{\text{total cell volume}}$$

$$\gamma = \frac{\frac{4}{3}\pi r^3}{\frac{4}{3}\pi r^3 + (L - 2r)\pi r^2} + \alpha\frac{(L-2r)\pi r^2}{\frac{4}{3}\pi r^3 + (L-2r)\pi r^2}$$

$$= \frac{4r + (3L - 6r)\alpha}{3L - 2r} \quad (13)$$

Note that $\frac{4r}{3L-2r} < \gamma < 1$ (note $L > 2r$) if the polar caps are always fully filled. When $\alpha = 1$, $\gamma$ equals 1 as well.

For the glucose-growth experiment (average birth length 1.6 µ$m$) we determined that $\alpha = 0.31$, which leads to $\gamma = 0.52$ (using the last equation), indicating around 48% of the cell volume is filled with DNA. This approximation is slightly lower compared to findings from the Jacobs-Wagner lab who report a nucleoid to cytoplasmic area ratio between roughly 0.5 an 0.6 for E.coli [10]. When we simulate cell growth on LB we obtain $\alpha = 0.66$, which leads to $\gamma = 0.75$, meaning about 25% of the cell filled with DNA (compared to a nucleocytoplasmic of roughly 0.38 found by Gray et al.[10]). This is in line with the idea that faster growing, larger cells will have a more homogenous spread of ribosomes throughout their cytoplasm. In the case of the larger LB cells (which are on average 6.72 $\mu m$ at the end of the cell cycle compared to 3.21 $\mu m$ for glucose), cells exhibit multiple areas of high ribosome fluorescence throughout the middle part of the cell as a opposed to solely in the caps. They show a zig-zag-like structure throughout the cell. Therefore, we would not expect to see a similarly distinguished growth-rate disturbance after birth for the LB cells (fig. S8), as ribosomes should be divided more evenly across daughter cells. This will be further explored in the next two sections.

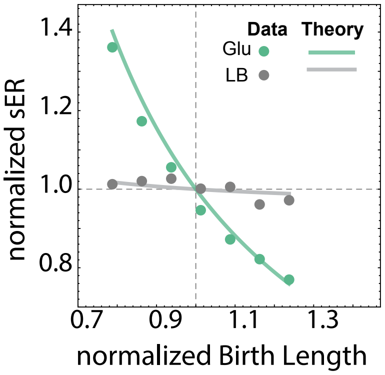


Figure S6: **Normalized birth growth rate (normalized age 0-0.05) for experimental E.coli data compared to a theoretical model line.** The alpha value representing the cylindrical part of the cell filled with ribosomes for the glucose theory line is 0, meaning assuming all ribosomes are found in the polar caps of the cell. For Lb the alpha value used is 0.9, meaning ribosomes are assumed to be spread more homogeneously throughout the cell.

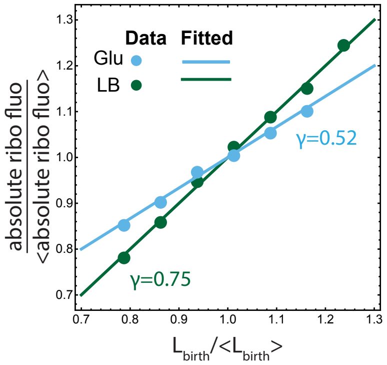


Figure S7: **Linear fit of normalized absolute ribosome fluorescence in a cell as function of its normalized Birth Length.** The fitted parameter is $\gamma$, representative of the fraction of cell's volume filled with ribosomes. This means 1-$\gamma$ gives the fraction of the cell part filled with nucleoid. For glucose the fitted $\gamma$= 0.52, for LB $\gamma$=0.75.

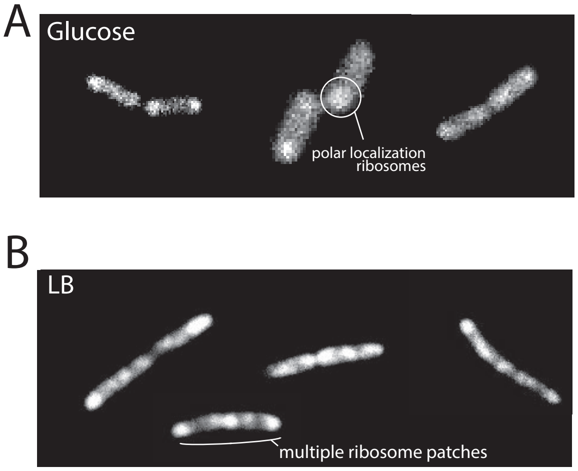


Figure S8: **Microscopy pictures of cells expressing mCherry-tagged L9 ribosomal proteins.** (A) For cells grown on glucose, ribosomes are primarily located in the polar caps of the cells. For cells grown on LB, ribosomes spread more throughout the cell's full length, in different patches due to the existence of multiple ORIs associated with fast growth.

##### S2.5.4 Homogenous spread of small proteins (incl. metabolic proteins) inside cells

Small proteins such as fluorescent proteins and metabolic proteins spread homogeneously through the cytosol[1,2], unlike large assemblies such as ribosomes which excluded from the midcell-positioned nucleoid. Uneven division will have no effect on the concentration of metabolic proteins in the newborn daughters. Regardless of the newborn size, daughter cells are expected to show approximately the same concentration of these proteins as their mother. This corresponds to a molecule with $\gamma = 1$. Indeed, as shown in fig. 3, the concentration of a constitutively expressed Lac protein does not seem to depend on birth size. This confirms our hypothesis that ribosomes are unevenly distributed between differently sized cells whilst metabolic protein concentration remains the same, creating the catabolic-anabolic imbalance underlying the growth rate variability.

##### S2.5.5 The concentration of ribosomes in single cells before and after division, with volume fraction $\gamma$ filled with ribosomes

Since the concentration of ribosomes of average-sized mother cells equals that of average-sized daughter cells, so $\langle c_B \rangle = \langle c_M \rangle$, we can use equation 12 to relate the ribosome concentration of a newborn cell with a certain $\alpha$ (or $\gamma$ value) to the concentration of ribosome of the average mother cell,

$$c_B = \frac{(4r - 3\langle L_M \rangle)(4\langle L_B \rangle r(2 - 3\alpha) + 3L_B \langle L_M \rangle \alpha)}{2\langle L_B \rangle (3L_D - 2r)(-3\langle L_M \rangle \alpha + 4r(3\alpha - 2))} \langle c_M \rangle. \tag{14}$$

(Note that for $\alpha = 1$ ( or $\gamma = 1$) we obtain $c_B = \langle c_M \rangle$, since the ribosomes are then distributed homogeneously throughout the cell and birth length no longer influences the concentration in new cells. So this applies for metabolic proteins but not ribosomal proteins.)

##### S2.5.6 Small daughter cells grow faster than large daughter cells after birth

Equation 2 gives the relation between the ribosome concentration and the growth rate from which we conclude that at cell birth, a daughter cell has growth rate,

$$\mu_B = k_r f_r c_{rB} \frac{1}{c_{tot}}, \tag{15}$$

and $c_B$ equals equation 14. When we consider, for simplicity, that all ribosomes are in the poles then

$$\mu_B = \frac{3\langle L_M \rangle - 4r}{6L_B - 4r} f_r k_r \langle c_{rM} \rangle \frac{1}{c_{tot}}, \tag{16}$$

which leads to

$$\frac{\mu_B}{\langle \mu_B \rangle} = \frac{\frac{3\langle L_M \rangle - 4r}{6L_B - 4r} f_r k_r \langle c_M \rangle}{\frac{3\langle L_M \rangle - 4r}{6\langle L_B \rangle - 4r} f_r k_r \langle c_M \rangle} = \frac{3\langle L_B \rangle - 2r}{3L_B - 2r} \tag{17}$$

This equation indicates that smaller-than-average daughter cells, $L_B < \langle L_B \rangle$, grow faster than average-sized daughter cells who, in turn, outgrow larger-than-average daughter cells.

We can compare this theoretical description of $\mu_B$ as function $L_B$ to our experimental data (fig. S6). This figure shows we are able to describe the birth growth rate differences in different nutrient environments by using varying $\gamma$ values. Note that the $\gamma$ value that best describes the glucose data equals 0 and that does not agree with our fitted value. This highlights that this simple phenomenological description is not able to capture all observed variation – we are still assuming that daughter cells derive from average mothers, which is apparently insufficient to explain the data. To improve the predictive power, we next describe a dynamic model of growth, ribosome regulation and uneven mother-cell division which we can run over thousands of consecutive cell cycles to address the importance of mother cells deviating from average mother cell behaviour.

#### S2.6 A deterministic model to describe the dynamics of birth-size-dependent, single-cell growth rates from birth to division

##### S2.6.1 Metabolism and protein expression model

Above we have described a model that aims to explain the experimentally-observed, growth-rate disturbance immediately after cell birth, assuming that all mothers resemble average mothers. We will now formulate a dynamic, deterministic model of growing cells from birth to division, followed by random, uneven cell-division and deterministic partitioning of molecular content. With this model we can simulate consecutive cell divisions and take deviating mother cells into account. This model includes compensation of the growth rate disturbance at division by control of metabolic and ribosomal protein expression by ppGpp and septum formation and invagination (to explain the enhanced length growth rate, which we observe at the end of the cell cycle in our experimental data (see Fig. 1 main text)).

Our model consists of the following differential equations, which describe the rate of change in the concentrations of amino acids ($a$), metabolic proteins ($m$), and ribosomes ($r$),

$$\begin{aligned}
\dot{c}_a &= k_m c_m \frac{1}{1+\left(\frac{c_a}{K_c}\right)^2} - k_r f_r(c_a) c_r, \\
\dot{c}_r &= \epsilon_r(c_p) \frac{k_r}{N_r} f_r(c_a) c_r - \mu_V c_r, \\
\dot{c}_m &= (1-\epsilon_r(c_p)) \frac{k_r}{N_m} f_r(c_a) c_r - \mu_V c_m. \qquad (18)
\end{aligned}$$

We use the following notation: $k_m$ represents the phenomenological catalytic rate constant of metabolism for amino acid synthesis, $c_m$ corresponds to the total concentration of metabolic proteins, $K_c$ is the inhibition constant of metabolism with respect to amino acids, $k_r$ is the catalytic rate of the ribosomes (in amino acids per second, $c_r$ equals the concentration of ribosomes, $f_r(c_a)$ equals the saturation of the ribosome with amino acids (modelled as $c_a/(c_a + K_r)$ with $K_r$ as a Michaelis-Menten constant), $N_r$ and $N_m$ represent the number of amino acids in ribosomes and average metabolic proteins, the specific volume growth rate is defined as $\mu_V$. The production of ribosomes and metabolic proteins depends on $\epsilon_r(c_p)$, which incorporates the regulation of protein expression by ppGpp with concentration $p$. Below we will derive an equation for $\epsilon_r(c_p)$. Below we will refer to $\left(1+\left(\frac{c_a}{K_c}\right)^2\right)^{-1}$ as $f_m(c_a)$.

##### S2.6.2 The growth-rate disturbance at cell birth is compensated for during the cell cycle by the regulatory action of ppGpp

Above we concluded smaller-than-average newborn cells grow faster than average newborns due to their higher-than-average ribosome concentrations, while larger-than-average newborns grow slower due to their lower-than-average ribosome concentrations. This occurred because the ribosome concentration is birth-size dependent, while the metabolic protein concentration is not. Figure 3 indicates that individual cells can adjust this ribosomal-metabolic protein-concentration imbalance via the regulatory action of ppGpp[17] such that cells with a birth size that deviates from the mean size steer themselves towards ribosome and metabolic-protein concentrations that resemble the average cell. In smaller-than-average cells the excess ribosome deplete the amino acid pool, such that its saturation with amino-acid-loaded tRNA reduces (its $f_r(c_a)$ drops relative to the average cell), which leads to the synthesis of ppGpp by RelA, which searches for ribosomes bound to unloaded tRNAs and then makes ppGpp. So, in smaller-than-average cells the concentration of ppGpp rises. In larger-than-average, the opposite occurs RelA finds less ribosomes bound to unloaded tRNAs ($f_r(c_a)$ is now higher than the average cell) leading to a lower ppGpp concentration because of a lower activity of RelA and the same activity of SpoT, the ppGpp degrading enzyme. So, in larger-than-average cells the concentration of ppGpp drops. Since binding of ppGpp to RNA polymerase lowers its affinity for ribosomal promoters and enhances its affinity for metabolic promoters, smaller-than-average cells restore their ribosome versus metabolic-protein imbalance by making relative more metabolic proteins while larger-than-average cells do so by making relatively more ribosomes. This continues until a steady state is reached which corresponds to the state of the average cell.

This mechanism is captured by the following extensions to the model shown in equation 18. Since the ppGpp regulatory system responds to excesses or shortages of amino acids, we did not incorporate tRNAs in our model, only amino acids. RelA synthesises ppGpp from GTP when uncharged tRNAs are bound tot the A-site of ribosomes[17]. Then $f_r(c_a)$ is low, such that the activity of RelA is proportional to $1 - f_r(c_a)$ with $f_r(c_a) = \frac{c_a}{K_r + c_a}$. SpoT is a hydrolase that converts ppGpp back into GTP. To design a system that steers the ribosome saturation back the desired level $f_o$ of average cells, we assume that RelA and SpoT are both saturated with their substrates, i.e. GTP for RelA and ppGpp for SpoT. Then we obtain for the differential equation for the concentration of ppGpp,

$$\begin{aligned}\dot{c}_p &= v_{RelA} - v_{SpoT} \\ &= k_{relA}(1 - f_r(c_a)) - k_{spoT} \\ &= k_{relA}\left(1 - \frac{k_{spoT}}{k_{relA}} - f_r(c_a)\right) \\ &= k_{relA}(f_o - f_r(c_a)).\end{aligned}$$

We defined the $f_o = 1 - \frac{k_{SpoT}}{k_{RelA}}$ which represents the steering direction of the ppGpp regulatory system since at steady state this differential equation leads to $f_r(c_a) \to f_o$ in all cells. This means that the steady state amino acid concentration always follows from,

$$f_o = \frac{c_{a,s}}{K_r + c_{a,s}} \quad \Rightarrow \quad c_{a,s} = \frac{f_o K_r}{1 - f_o}.$$

We choose a value of $f_o = 0.8$ which reflects experimental data[4]. The steering of $f_r(c_a) \to f_o$ is done by the adjustment of the ribosome and metabolic-protein concentration via ppGpp-mediated regulation of protein expression. This is captured by the term $\epsilon_r(c_p)$, which represent the influence of ppGpp on the protein synthesis rate of ribosomes, while $1 - \epsilon_r(c_p)$ captures ppGpp's influence on the synthesis rate of metabolic proteins,

$$\epsilon_r(c_p) = \frac{K_p}{K_p + c_p}, \quad \Rightarrow \quad 1 - \epsilon_r(c_p) = \frac{c_p}{K_p + c_p}, \tag{19}$$

indicating that ppGpp activates metabolic-protein synthesis and inhibits ribosome synthesis.

##### S2.6.3 Model design requirements

A requirement of the model is that it agrees with several fundamental aspects of *E.coli* growth and its regulation,

1. the relationship between the steady-state ribosomal protein fraction $\phi_r = \frac{c_r}{c_r + c_m}$ and the growth rate is linear[7,20]. The model agrees with this, because (given equation 18),

$$\dot{c}_r + \dot{c}_m = \left(\epsilon_r \frac{1}{N_r} + (1 - \epsilon_r)\frac{1}{N_m}\right) k_r f_r(c_a) c_r - \mu_v (c_r + c_m)$$

   such that at steady state,

$$\mu_v = \left(\epsilon_r \frac{1}{N_r} + (1 - \epsilon_r)\frac{1}{N_m}\right) k_r f_o \phi_r \quad \Rightarrow \quad \phi_r = \frac{\mu_v}{k_r' f_o}. \tag{20}$$

   The last relationship is proportional and not linear in $\mu_v$ because we do not consider inactive ribosomes. Adding this would not change the model, so we left this out.

2. The observed relationship between the steady-state ppGpp concentration and the growth rate appears proportional to $1/\mu_V$ [12,14]. Our model also has this characteristic too. The steady state of the amino acid concentration $\dot{c}_a = 0$ leads to,

$$\frac{c_m}{c_r} = \frac{k_r f_o}{k_m f_m(c_{a,s})}$$

from $\dot{c}_m = 0$ and $\dot{c}_r = 0$ we obtain,

$$c_m = (1 - \epsilon(p)) \frac{\frac{k_r}{N_m} f_o c_r}{\mu_V}, \; c_r = \epsilon(p) \frac{\frac{k_r}{N_r} f_o c_r}{\mu_V},$$

such that (using equation 19)

$$\frac{c_m}{c_r} = \frac{k_r f_o}{k_m f_m(c_{a,s})} = \frac{1 - \epsilon(p)}{\epsilon(p)} \frac{N_r}{N_m} = \frac{c_p}{K_p} \frac{N_r}{N_m} \tag{21}$$

and since

$$\phi_r = \frac{c_r}{c_r + c_m} = \frac{1}{1 + \frac{c_m}{c_r}} \quad \Rightarrow \quad \frac{c_m}{c_r} = \frac{1}{\phi_r} - 1 = \frac{k'_r f_o}{\mu_V} - 1$$

where we used equation 20. Finally this leads to,

$$c_p = \frac{K_p N_m}{N_r} \left( \frac{k'_r f_o}{\mu_V} - 1 \right), \tag{22}$$

in agreement with the trend of the experimental findings.

3. the relationship between the nutrient concentration and growth rate is hyperbolic (Monod's relation). The model shows this too, consider equation 20 and 21, which together lead to,

$$\mu_V = \left( \frac{1}{N_r} + \frac{1}{N_m} \right) k_r f_o \frac{k_m f_m(c_{a,s})}{k_r f_o + k_m f_m(c_{a,s})},$$

when we assume that metabolism saturates with the nutrient concentration, $s$, as,

$$f_m(c_{a,s}) = \frac{\frac{s}{K_s}}{1 + \frac{s}{K_s} + \frac{c_{a,s}}{K_a}},$$

then the growth rate can be written as,

$$\mu_V = \mu_{max} \frac{s}{K_M + s}, \quad \text{with: } \mu_{max} = \left( \frac{1}{N_r} + \frac{1}{N_m} \right) \frac{k_r f_o K_a k_m}{K_a(k_m + k_r f_o)}, \; K_M = \frac{f_o(c_{a,s} + K_a) k_r K_s}{K_a(k_m + f_o K_r)},$$

which agrees with Monod's relation.

###### S2.6.4 Including two different types of length growth makes the model comparable to the experimental data

We also desired that the model could explain the increase in the length growth rate close to the end of the cell cycle. This observation can be explained by cap growth during the end of the cell cycle. The septum then forms by invagination such that the diameter of the growing (midcell) cell part becomes smaller and if the volumetric growth rate remains constant the length growth rate has to increase due to the ever smaller cell diameter (Fig. S10).

We define two growth rates, one accounts for cell growth during the period before septum formation (invagination), so growth at a constant cell radius, and the other for cell growth during the period when the septum is being formed and the diameter becomes smaller in midcell region until the cell divides.

1. **Cell growth by growth of the cylindrical region.** The length growth rate at a constant diameter, so before septum formation, can be obtained from the following relations.

   We can rewrite equation (4) to express the length of a cell in terms of its volume,

$$L(V) = \frac{V + \frac{2}{3}\pi r^3}{\pi r^2}.$$

   The specific growth rate of the length of a single cell is related to the specific growth rate of its volume as (see equation 5),

$$\mu_L = \frac{1}{L}\dot{L} = \frac{V}{L}\frac{\partial L}{\partial V}\mu_V = \frac{\pi r^2(L-2r) + \frac{4}{3}\pi r^3}{L}\frac{1}{\pi r^2}\mu_V = \left(1 - \frac{2}{3}\frac{r}{L}\right)\mu_V, \tag{23}$$

   indicating that the length growth rate is not constant during balanced growth when $\mu_V$ is constant, due to is dependency on $L$.

2. **Cell growth by midcell cap (septum) formation** During the last phase of the cell cycle *E. coli* cells form two new caps (due to midcell invagination) before they divide. This results in a significantly different relation of specific length growth to specific volume growth of the cell.

   Because two new caps are formed simultaneously, the growth can be visualised as a growing sphere. Imagine a complete sphere and a vertical plane cutting through its centre coordinate. Left and right of the plane we have half a sphere. Septum formation is the formation of those two half spheres as function of time starting from the plane, so we make discs with a reducing diameter.

   We define the distance of this growing sphere from the plane as $L_{cap}$, which ranges from $\frac{-L_{cap}}{2}$ to $\frac{L_{cap}}{2}$. Note that $0 \le L_{caps} \le 2r$. We find the volume of both growing caps after growth to distance $L_{cap}$ from,

$$V_{caps}(L_{cap}) = \int_{\frac{-L_{cap}}{2}}^{\frac{L_{cap}}{2}} \int_{-\sqrt{r^2-x^2}}^{\sqrt{r^2-x^2}} \int_{-\sqrt{r^2-x^2-y^2}}^{\sqrt{r^2-x^2-y^2}} dzdydx = r^2 L_{cap}\pi - \frac{1}{12}L_{cap}^3\pi, \tag{24}$$

   (note this last equation with $L_{cap} = 2r$ gives the volume of a sphere $\frac{4}{3}\pi r^3$). This last equation gives,

$$\frac{\partial V_{caps}}{\partial L_{cap}} = r^2\pi - \frac{L_{cap}^2}{4}\pi.$$

During cap growth, we have a $\mu_V$ that results from protein synthesis and this related to length growth during cap formation as (note growth in the cell length $L$ equals growth in $L_{cap}$ when only the caps grow),

$$\mu_V = \frac{1}{V}\frac{\partial V_{caps}}{\partial L_{caps}}\frac{dL_{caps}}{dt} = \frac{1}{V}\frac{\partial V_{caps}}{\partial L_{caps}}\frac{dL}{dt} = \frac{L}{V}\frac{\partial V_{caps}}{\partial L_{caps}}\mu_L = \frac{L}{V}\left(r^2\pi - \frac{L_{caps}^2\pi}{4}\right)\mu_L,$$

which can be rewritten to express the length growth rate as a function of current volume $V$, length $L$ and the volume growth rate as,

$$\mu_L = \frac{V}{L\left(r^2\pi - \frac{L_{caps}^2\pi}{4}\right)}\mu_V. \tag{25}$$

This last equation gives, with $V = V_{cyl} + V_{caps} = \pi r^2 L_{cyl} + r^2 L_{cap}\pi - \frac{1}{12}L_{cap}^3\pi$ and $L = L_{cyl} + L_{cap}$, indeed a function $\mu_L/\mu_V$ that rises almost exponentially with $L_{cap}$ (note: $0 \le L_{cap} \le 2r$, realistic values $r = 0.6\ \mu m$, $L_{cyl} = 1.2\ \mu m$) as we see in the data. Note that $\mu_L/\mu_V = 1$ when $L_{cap} = 0$ as expected.

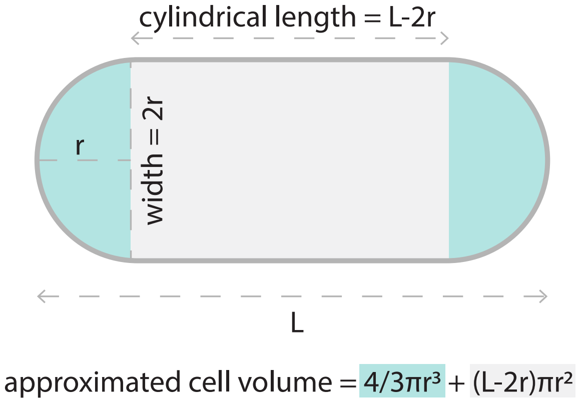


Figure S9: **Blueprint of the cell shape of a rod shaped bacterium, assuming cells are a perfect cylinder with a hemispherical cap on each end.** This shows how the length of one pole (its radius) equals half of the cell width. Then, the length of the cylindrical part of the cell is L-2r where L is the leng of the whole cell.Taking this into account the volume of rod shaped cells can be estimated to their polar volume in addition to their cylindrical value: $\frac{4}{3}\pi r^3 + (L-2r)\pi r^2$. Of course, in reality cells will not portray such perfectly shaped rods, but this serves as the blueprint for how we consider cell shape in our model and data analyses.

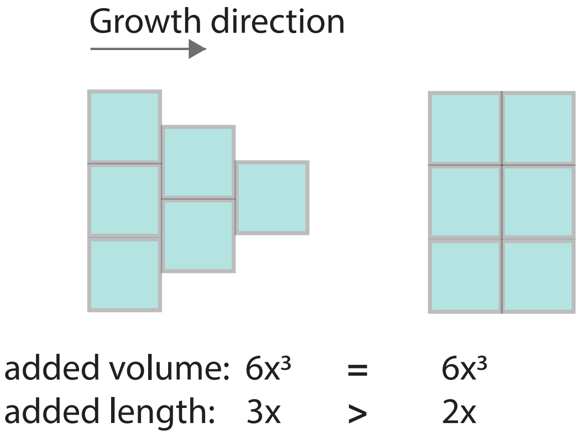


Figure S10: **The shape of a growing structure will affect the volumetric versus length growth rate.** As illustrated in a simplified fashion here, when adding the same volume (biomass) while forming a cap (approximated on the left) there will be a larger length increase compared to forming a cylinder (approximated on the right). Therefore, a constant volumetric growth rate will show an increasing length growth rate when growing spherical cap, which becomes smaller towards its ends.

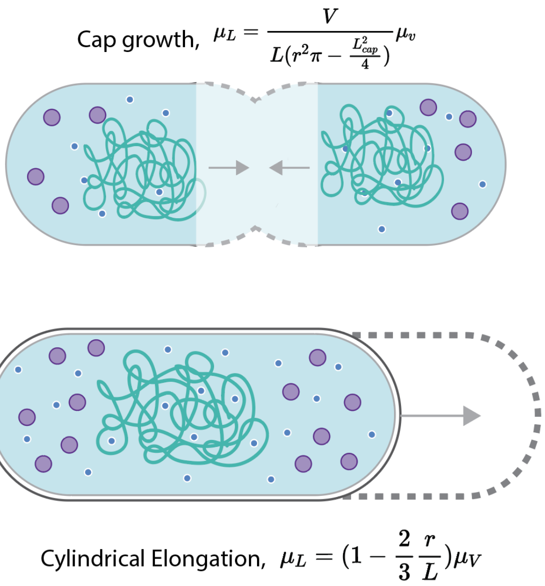


Figure S11: **Illustration of the two considered types of cell length growth ($\mu_L$), and their relation to the specific volume growth rate ($\mu_V$).** Cylindrical elongation and cap formation result in two different functions for $\mu_L$. Since the growth rate measured in the microscopy data depends on length data, not volume, this distinction between might influence measured results.

##### S2.6.5 Timing and initiation of two growth modes (cylindrical and midcell caps)

We assume that during the last phase of the cell cycle a fraction $\beta$ of synthesised volume is allocated to cylindrical (region) growth and the remaining $1-\beta$ to cap formation. This $\beta$ depends on the growth rate[6], on which we will elaborate later on. Before the cell starts making its midcell caps, all volume synthesis is cylindrical growth. We assume that at the point of division, where the mother cell resembles two touching rod-shaped daughter cells, the newly formed caps are still connected (their contact volumes form a small disk). Thus, the two caps are together not a complete sphere, but have length $r$ minus the length of a spherical segment $\delta r$. Therefore, the volume of the two newly formed caps can be obtained from substituted $L_{cap} = 2r - 2\delta r$ in equation 24 giving $\frac{4}{3}\pi r^3 - 2\pi r\delta r^2 + \frac{2}{3}\pi\delta r^2$.

The total volume added during the cap-formation phase is,

$$\begin{aligned}\Delta V_{tot} &= \Delta V_{cap} + \Delta V_{cyl} \\ &= \beta\Delta V_{tot} + (1-\beta)\Delta V_{tot} \\ &= \frac{4}{3}\pi r^3 - 2\pi r\delta r^2 + \frac{2}{3}\pi\delta r^2 + (1-\beta)\Delta V_{tot} \\ &= \frac{1}{\beta}\left(\frac{4}{3}\pi r^3 - 2\pi r\delta r^2 + \frac{2}{3}\pi\delta r^2\right)\end{aligned} \tag{26}$$

We know about the volume added as cylinder that,

$$\Delta V_{cyl} = (1-\beta)\Delta V_{tot}$$

Substituting (26) into the above gives,

$$\Delta V_{cyl} = (1-\beta)\frac{1}{\beta}\left(\frac{4}{3}\pi r^3 - 2\pi r\delta r^2 + \frac{2}{3}\pi\delta r^2\right) = \pi r^2 L_{CylTot}$$

where $L_{CylTot}$ is the length added during the last phase to the cylindrical part.

From the last equation we can determine $L_{CylTot}$, the total added cylindrical length in the last phase. We can add to this length the total length added during cap growth,

$$\Delta L_{\text{2nd phase}} = L_{CylTot} + L_{CapTot} = \frac{(1-\beta)\frac{1}{\beta}\left(\frac{4}{3}\pi r^3 - 2\pi r\delta r^2 + \frac{2}{3}\pi\delta r^2\right)}{\pi r^2} + 2r - 2\delta r$$

Thus, we divided one cell cycle into two distinct growth phases: during the first growth phase starting right after birth all the growth in volume contributed to cylindrical growth. During the second phase, a fraction $\beta$ of the added volume goes into forming the new caps, the rest $1-\beta$ is invested in cylindrical growth. To ensure the cell adds $\Delta L$ during its entire cell cycle, the second growth phase is initiated when the cell's length equals,

$$\Delta L_{\text{1st phase}} = \Delta L - \Delta L_{\text{2nd phase}}$$

At the timepoint when the cell meets this condition, the cell grows cylindrical and invaginates and the change in cellular length can now be described as,

$$\begin{aligned}\dot{L} &= \dot{L}_{cylinder} + \dot{L}_{caps} \\ &= \frac{\partial L_{cyl}}{\partial V_{cyl}} \dot{V}_{cyl} + \frac{\partial L_{cap}}{\partial V_{cap}} \dot{V}_{cap} \\ &= \frac{1}{r^2 \pi}(1-\beta)\mu_V V + \frac{1}{r^2\pi - \frac{L_{cap}^2 \pi}{4}} \beta \mu_V V,\end{aligned} \tag{27}$$

with $V_{cyl}(L_{cyl}) = \pi r^2 L_{cyl}$ and $V_{cap}(L_{cap})$ is given by equation 24.

Finally, we run the model from cell birth to division, where its $\Delta L$ is determined from equation (6). The birth lengths are random variables chosen in accordance with experimental data of birth length conditional on mother size (fig. S12). So given the length of her mother, the daughter cells gets a particular starting length.

#### S2.7 Simulating thousands of cell generations to capture the non-steady state mother effect

Since, mother cells, perturbed at birth, do not fully compensate this perturbation prior to division (figs. S3, S4 and S13), the metabolic state of newborn daughter depends on their mother cell state at the time of division. To capture how this mother-effect influences our data (illustrated in fig. 3C main text), we simulated thousands of consecutive cell cycles (fig. S12). Each time a cell has added $\Delta L$, and will thus divide, the $L_{birth}$ of the next daughter cell will be sampled from the experimentally fitted bivariate normal distribution given mother length $L_{division}$ (fig. S12). This new length, combined with the molecular concentrations of the mother, determines the new daughter's molecular state.

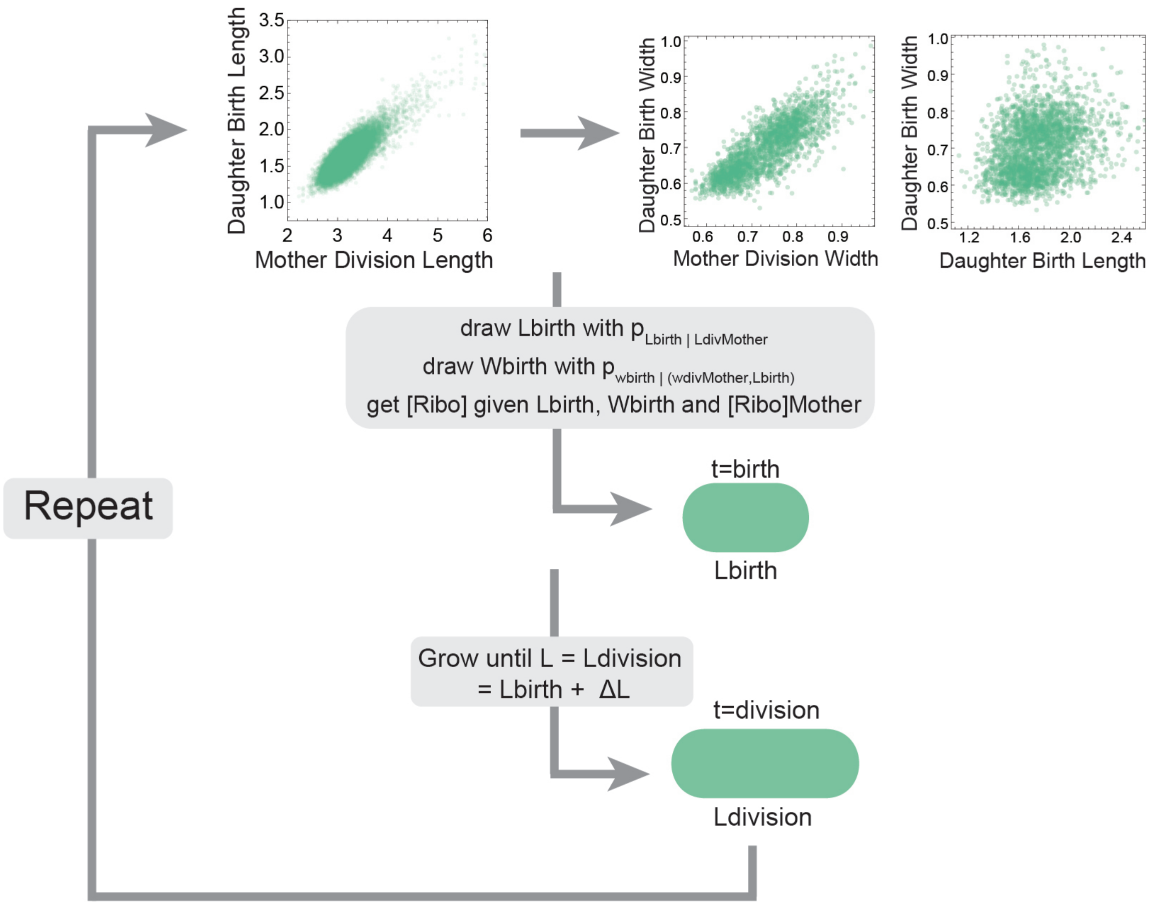


Figure S12: **The birth length and width of daughter cells of mothers with different division sizes, from our experimental data.** The corresponding fitted multinomial distributions were used in the model simulation to draw birth lengths and widths corresponding to a probability distribution given the simulated mother's length and width.

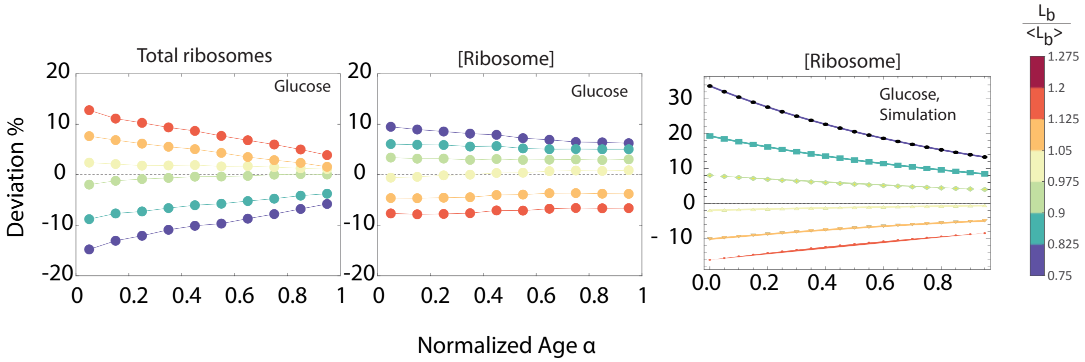


Figure S13: **Comparison of experimental and simulated ribosome content deviations**. Shown is a comparison of the simulated (right panel) ribosome concentration deviation from average over the whole cell cycle, for all the different birth-length bins, to that of the experimental data (middle panel), and the experimental absolute (total) ribosome deviation (left panel).

Table S4: Model parameters

| Parameter | Unit | Description | Values | | |
|---|---|---|---|---|---|
| | | | Glc | LB | Literature |
| kr | $min^{-1}$ | $k_{cat}$ of protein synthesis by ribosomes | 1320 | 1320 | 1320[5] |
| kc | $min^{-1}$ | Phenomenological "efficiency" of metabolic enzymes, interpreted as the enzymatic rate converting current nutrient source into cellular precursors (here simply amino acids). Seen as $k_{cat}$ of amino acid synthesis by metabolic enzymes, their saturation by nutrients and an efficiency measure. | 25 | 50 | avg enzymatic $k_{cat}$ = 100[3] |
| Kr | $\mu$M | $K_m$ of ribosomes for charged tRNA (here, amino acids) | 80 | 80 | ? |
| Kc | $\mu$M | Inhibition constant for amino acid synthesis by metabolic enzymes | 800 | 800 | 100[8] |
| Nm | - | Number of amino acids in one metabolic enzyme | 400 | 400 | ~350[25] |
| Nr | - | Number of amino acids in one ribosome | 1500 | 1500 | 7336[5] |
| $\beta$ | - | Fraction of volume growth rate attributed to cap-growth during end of the cell cycle. $\beta$=1 implies all volume growth is attributed to forming new caps. | 0.6 | 0.35 | – |
| kRela | $min^{-1}$ | $k_{cat}$ for ppGpp synthesis by RelA | 3000 | 3000[11] | |
| $f_o$ | - | Optimal saturation of ribosomes with amino acids. Setpoint for the ppGpp-steered integral control system. | 0.725 | 0.725 | 0.8[5] |
| $L_M$ | $\mu$m | Cell length of average mother at division | 3.51 | 6.62 | * |
| $L_0$ | $\mu$m | Birth length of an average daughter cell | 1.76 | 3.31 | * |
| d | $\mu$m | Radius of the cell. Length of one cap. | 0.25 | 0.5 | * |
| $\alpha$ | - | Fraction of the cylindrical part of a cell filled with ribosomes, where $\alpha$=0 implies ribosomes are only found in the caps. | 0 | 0.9 | |
| Kp | $\mu$M | Concentration of ppGpp at which half of the ribosomes are attributed to synthesize ribosomes | 5000 | 5000 | ? |

* These values have been extracted from the experimental data. Average mother length is taken directly from the experimental data, the average daughter length is chosen as $L_M/2$, since there is a short time delay in the microscopy data which makes it seem as if the average daughter is a little larger than half the average mother. Since the model simulates length growth continuously and does not incorporate intermediary time-steps, using the average experimental daughter length would give a faulty representation of division with a tendency to divide larger.
